# Exploiting evolutionary herding to control drug resistance in cancer

**DOI:** 10.1101/566950

**Authors:** Ahmet Acar, Daniel Nichol, Javier Fernandez-Mateos, George D. Cresswell, Iros Barozzi, Sung Pil Hong, Inmaculada Spiteri, Mark Stubbs, Rosemary Burke, Adam Stewart, Georgios Vlachogiannis, Carlo C. Maley, Luca Magnani, Nicola Valeri, Udai Banerji, Andrea Sottoriva

**Author notes:** equal contribution (AA: wet lab work, DN: dry lab work).

## Abstract

Drug resistance mediated by clonal evolution is arguably the biggest problem in cancer therapy today. However, evolving resistance to one drug may come at a cost of decreased growth rate or increased sensitivity to another drug due to evolutionary trade-offs. This weakness can be exploited in the clinic using an approach called ‘evolutionary herding’ that aims at controlling the tumour cell population to delay or prevent resistance. However, recapitulating cancer evolutionary dynamics experimentally remains challenging. Here we present a novel approach for evolutionary herding based on a combination of single-cell barcoding, very large populations of 10^8^–10^9^ cells grown without re-plating, longitudinal non-destructive monitoring of cancer clones, and mathematical modelling of tumour evolution. We demonstrate evolutionary herding in non-small cell lung cancer, showing that herding allows shifting the clonal composition of a tumour in our favour, leading to collateral drug sensitivity and proliferative fitness costs. Through genomic analysis and single-cell sequencing, we were also able to determine the mechanisms that drive such evolved sensitivity. Our approach allows modelling evolutionary trade-offs experimentally to test patient-specific evolutionary herding strategies that can potentially be translated into the clinic to control treatment resistance.

## Introduction

Although targeted cancer therapies are effective in many patients (Zhang et al., 2009), their efficacy is impeded by treatment resistance, currently an intractable problem in cancer. Resistance is often mediated by redundancies in downstream signalling pathways (Holohan et al., 2013), cell phenotypic plasticity (Meacham and Morrison, 2013), and most importantly, intra-tumour heterogeneity (ITH) (McGranahan and Swanton, 2015). The high level of ITH in the majority of cancers (McGranahan and Swanton, 2017) implies that pre-existing cancer subclones that are drug resistant because of heritable genetic (Pao et al., 2005) or epigenetic (Shaffer et al., 2017; Sharma et al., 2010) alterations are invariably present when treatment starts (Diaz et al., 2012; Misale et al., 2012), thus leading to Darwinian adaptation (Greaves and Maley, 2012). In addition, drug-tolerant cancer cells or ‘persistors’ can survive and acquire *de novo* heritable alterations that give rise to fully resistant subclones during or after treatment (Hata et al., 2016; Nichol et al., 2016). The emergence of pre-existing populations that prior to treatment are fitness neutral (or even deleterious) (Hall et al., 2009) and are positively selected by intervention can be recapitulated in the lab, as first demonstrated by the classical Luria-Delbruck experiment in bacteria (Luria and Delbrück).

This implies that cancers are unlikely to be successfully treated with a single agent, as we often observed in the clinic (Gillies et al., 2012). Whereas combination strategies are often highly toxic and impractical, relatively little is known about the most effective sequence of agents. Administering a drug can sensitise cancer cells to a second drug, a phenomenon known as *collateral sensitivity*, which has been demonstrated experimentally in seminal studies in bacteria (Imamovic and Sommer, 2013; Nichol et al., 2015; Pál et al., 2015), malaria (Kirkman et al., 2018) and cancer (Hall et al., 2009; Zhao et al., 2016b). This is based on the observation that in evolution, adaptations frequently incur a cost. As in ecological systems, developing a new trait such as resistance to cancer treatment likely comes at the expense of other features, such as loss of adaptive response to other stimuli (Gatenby et al., 2009; Merlo et al., 2006), leading to ‘evolutionary trade-offs’ (Fuentes-Hernandez et al., 2015). Cost of resistance has been observed in distinct pathogenic organisms (Hughes and Andersson, 2015a) as well as in cancer (Siravegna et al., 2015).

Evolutionary herding aims at exploiting trade-offs to control tumour evolution in our favour. The goal is directing the evolution of the tumour population using Darwinian adaptation to a drug. When a second drug is administered, the clonal composition of the population is different from the start, and this can lead to increased sensitivity, or even complete extinction (Pluchino et al., 2012; Zhao et al., 2016a). In this scenario, because evolutionary herding has changed the clonal structure of the population, collateral drug sensitivity is likely to be persistent rather than transient and can be exploited as a form of synthetic lethality. Therapeutic strategies that rely on deterministic evolutionary herding and controlled cancer evolution are also less subject to stochastic temporary effects and cell plasticity, and hence more likely to be effective in the clinic.

Current experimental approaches are inadequate to study evolutionary herding because they are limited to small populations that do not recapitulate the extensive intra-tumour heterogeneity present in human malignancies. Moreover, current approaches introduce substantial biases due to short timescales and re-plating, and rely on escalating drug doses that preferentially select for *de novo* evolution of small-effect alleles at multiple loci, rather than *pre-existing* highly resistant subclones (Nichol et al., 2017). Cell plasticity and drug tolerance, instead of Darwinian adaptation, often occurs in current model systems, leading to resistance that is non-heritable, potentially reversible, and that does not represent what happens in the clinic. Non-heritable drug resistance can arise through epithelial-mesenchymal transition (Shibue and Weinberg, 2017) or upregulation of drug-efflux pumps (Gottesman and Pastan, 1993). Although these are very important cellular mechanisms of resistance, they do not pertain to clonal evolution, which drives persistent resistance in human cancers over long timescales.

Here, we present a novel experimental approach to study evolutionary herding quantitatively and demonstrate the evolutionary determinants of collateral drug sensitivity by clonal herding of cancer cell populations.

## Results

### Evolutionary herding of resistant cells through fitness landscapes

The relationship between heritable information, whether genetic or epigenetic, and the corresponding cellular phenotype, can be represented by the classical *fitness landscape* model (Wright). Cellular phenotypes are multifaceted and arise as a product of the complex interactions between heritable factors and the environment. If we summarise the fitness of these complex phenotypes with respect to a certain condition or environment by a single value, the genotype-phenotype relationship can be represented as an *n+1* dimensional space whereby the alleles present at *n* (epi)genetic loci are mapped to the relative fitness advantage they confer. A single cell can therefore be represented by a point in this landscape corresponding to its (epi)genetic state. As populations proliferate and randomly mutate, cell lineages move around the landscape. In a simple illustrative drug-free scenario (Figure 1A), multiple cells, each characterised by a certain genotype (*x*, *y* and *z*), are scattered around a neutral ‘flat’ fitness landscape because of genotypic mutations. When a drug is applied (e.g. drug 1), the fitness landscape changes, and genotypes that were previously neutral (or even slightly deleterious) may become advantageous under the new condition (e.g. *y* and *z*), and outcompete the rest (e.g. *x*). Due to Darwinian selection, populations in lower fitness elevations will likely go extinct, whereas populations in fitness ‘peaks’ will prosper. This makes populations appear to ‘climb’ higher and higher fitness peaks, leading to evolutionary adaptation.

**Figure 1:**
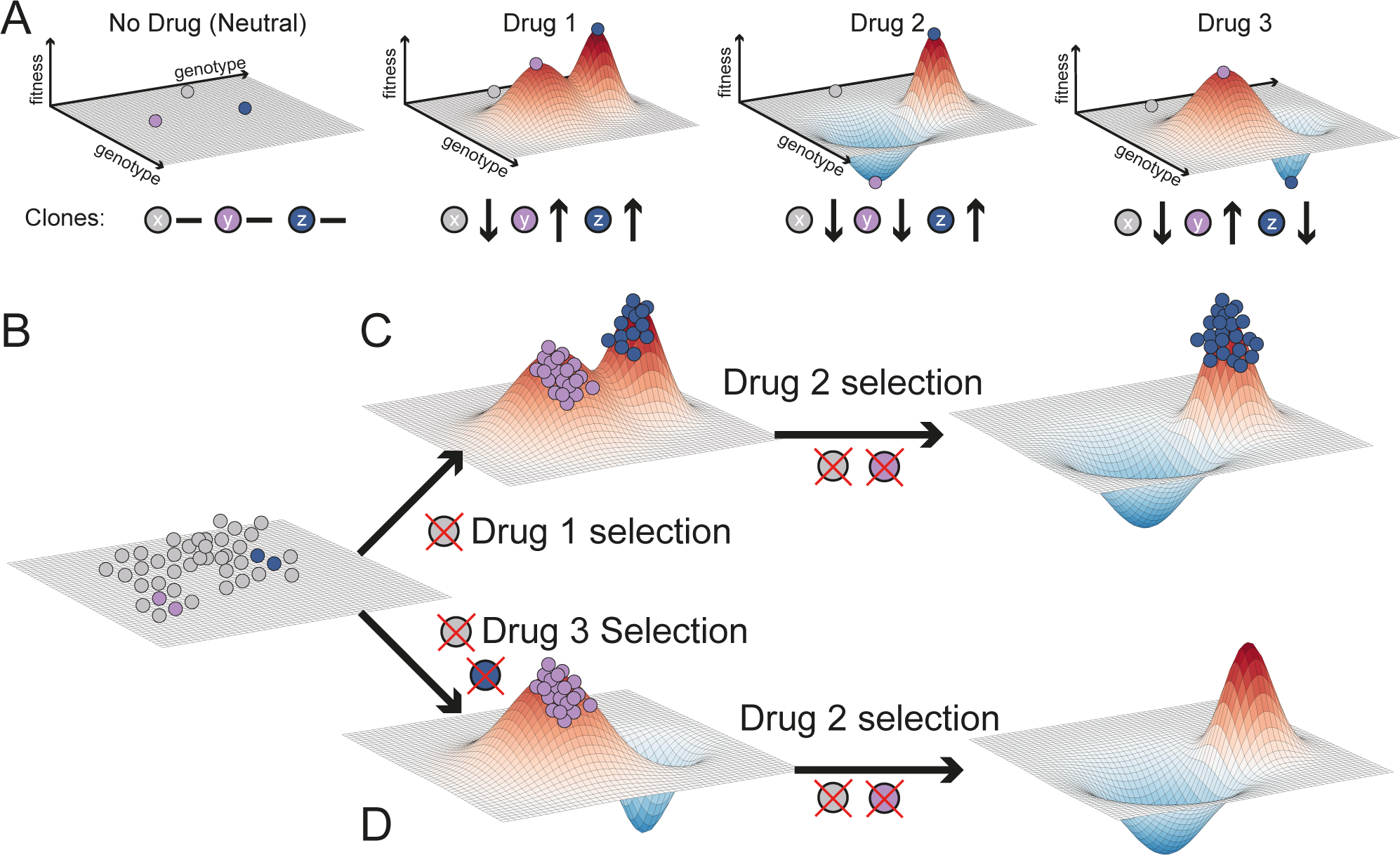
Evolutionary herding through fitness landscapes. **(A)** The selective effect of a drug on a heterogeneous population can be visualised as a fitness landscape. Genetically distinct cells are represented by points in the x-y plane, whilst their fitness within a certain environment is the z-value. Different drugs have different landscapes, selecting against different clones. For simplicity, here we assume in the absence of drug all clones to be equally fit (flat landscape). Drug 1 changes the landscape, selecting for y and z but against x. Drug 2 selects only for z and drug 3 only for y. **(B)** First, a population of cells is present at baseline, represented here as equally fit for simplicity. **(C)** Drug 1 selects clones that are resistant to drug 2. **(D)** Applying first drug 3 leads to evolutionary herding of a population that is entirely sensitive to drug 2. Here genotypes with values below the plain have negative fitness and so their frequency will decrease until they go extinct.

Different drugs may select for distinct phenotypes (e.g. *y* and *z* are differentially selected by drug 2 and 3 – Figure 1A). Using drugs with divergent fitness landscapes is the central idea of evolutionary herding. This concept is illustrated in Figure 1B. Tumourigenesis gives rise to a heterogeneous population of cancer cells that is the substrate for Darwinian selection to operate. When drug 1 is applied (Figure 1C), only populations that are around the new fitness peaks survive, while drug sensitive cells in fitness valleys go extinct. If then we expose the population to drug 2, which has an overlapping fitness peak, we select for a doubly resistant phenotype *z*, against which both drug 1 and 2 are ineffective. At this point we would have lost control of the tumour. Instead, if we first apply drug 3 (Figure 1D) this leads to selection for phenotype *y*. Because drug 2 shows differential fitness peaks with respect to drug 3, the sequence drug3-drug2 leads to an evolutionary trap in which the cancer cell population goes extinct (Zhao et al., 2016a). This is the principle of evolutionary herding that can be exploited to delay and potentially control drug resistance, thus significantly extending patient survival.

### Evolving resistance in large populations without re-plating

We first demonstrate evolutionary herding *in vitro* using the HCC827 non-small cell lung cancer line. HCC827 is an *EGFR* exon19del mutant lung cancer cell line sensitive to EGFR inhibition (Engelman et al., 2007). We used two small molecule inhibitors for herding: gefitinib, an EGFR inhibitor, and trametinib, a MEK1/2 inhibitor. To recapitulate the evolutionary dynamics of large populations, we employed a HYPERflask® cell culture system, wherein each flask has a capacity of up to 150 million cells (Figure 2A). To track clonal evolution we employed high complexity lentiviral barcoding (Bhang et al., 2015). By barcoding the cells at baseline and splitting them into distinct replicates (Figure 2B), we could determine whether resistant clones were pre-existing if the same barcodes were enriched post-treatment in different replicas. We first barcoded a population of one million cells with one million distinct barcodes, and then expanded it to ∼120M in a HYPERflask (Material and Methods). We call this initial baseline population the “POT” (Figure 2B). For each of the two drugs we seeded three HYPERflask replicates in addition to two HYPERflask as DMSO controls. Each HYPERflask was seeded with approximately 15 million cells from the same POT (i.e. most barcodes are common to all flasks) and expanded to 80-90% confluence. Thus, we achieved a total population of 120Mx3∼0.4 billion cells per drug arm (Figure 2B, Material and Methods).

**Figure 2:**
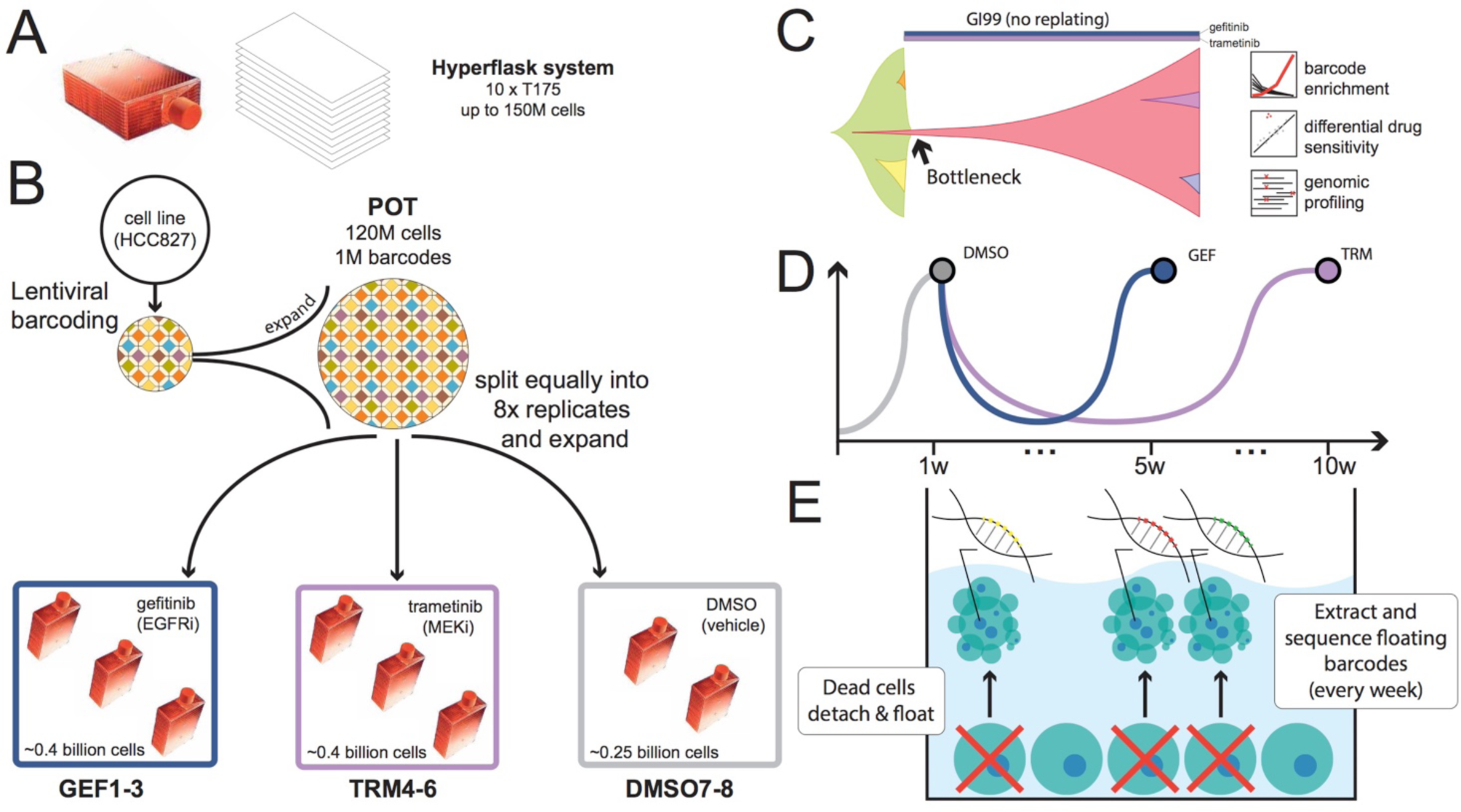
Experimental design. **(A)** The Corning® High Yield PERformance Flasks (HYPERflask®) cell culture vessel is a 10 layer 1720 cm^2^ total growth area system with polystyrene gas permeable surface that can reach the order of 150 million cells. **(B)** One million lung cancer cell line HCC827 cells were lentivirally barcoded before being expanded to a population (POT) of ∼120 million cells on a HYPERflask. Eight replicates were seeded with ∼12 million cells each and expanded to 120 million. The remaining POT cells are frozen for subsequent analysis. Of the eight seeded replicates, three are exposed to GI99 doses of gefitinib (GEF1-3), three to GI99 of trametinib (TRM4-6) and two are harvested immediately as controls (DMSO7-8). **(C)** Clonal evolution of a large population containing pre-existing resistant subclones that is exposed to high drug concentration (GI99) without re-plating. As in patients, a clonal bottleneck occurs by means of Darwinian selection for drug resistance. Barcode enrichment analysis, genomic profiling and drug screening is performed on the resistant population. **(D)** Schematic growth curves for gefitinib (4 weeks for resistant population to regrow) and trametinib (9 weeks for the resistant population to regrow). Media and drug are changed weekly. **(E)** When cells die, they detach and float in the media. At each media change (once per week), supernatant cells are harvested from the spent media and their DNA extracted for barcodes analysis and non-destructive tracking of tumour evolution.

These large populations allowed us to expose the cells to high drug concentrations without causing extinction and without the need for re-plating. This is because large populations are highly heterogeneous and likely to contain pre-existing resistant subclones that would survive high-dose drug exposure. We used GI99 concentrations (99% Growth Inhibition) until resistant clones grew back (Figure 2C and S1). Three HYPERflasks were drugged with gefitinib (40nM) and three with trametinib (100nM). Drug exposure in the gefitinib treated lines GEF1-GEF3, induced extensive cell death, thus causing a major population bottleneck (Figure 2D). Under constant drug concentration, the resistant population grew back and reached confluence again in 4 weeks. Drug exposure in the trametinib treated lines TRM4-TRM6 also induced extensive cell death and a resistant population grew back to confluence in 9 weeks (Figure 2D).

We reasoned that not only the surviving resistant cells at the end of the experiments were important for the analysis, but also that the cells that died during the experiment could prove informative on the temporal dynamics of the system. The idea is that the sum of the surviving cells attached to the plate and the dead cells floating in the media would contain information on the whole evolutionary history of the cell population. Moreover, we hypothesised that dead cells may be a representative sample of the live population and, like circulating tumour DNA in cancer patients (Domínguez-Vigil et al., 2018), could be used to monitor the temporal dynamics of the system non-destructively. Once a week at each media change, we collected the floating (dead) cells as pellets to extract DNA and perform barcode analysis (Figure 2E, Material and Methods).

We compared baseline (POT) vs resistant lines and confirmed decreased drug sensitivity for both gefitinib (Figure 3A) and trametinib (Figure 3B). To identify possible genetic mechanisms of resistance, we performed whole-exome sequencing at median 160x depth. We found a focal amplification of *MET* in gefitinib resistant lines (Figure 3C), consistent with previous results (*37*), that was confirmed by ddPCR (Figure S2). No amplification of *MET* was detected in trametinib resistant lines, suggesting that *MET* amplified subclones are gefitinib-resistant but may be trametinib-sensitive. The trametinib resistant lines shared a gain of chr1p and deletions in chr9, encompassing *CDKN2A* (Figure 3C, S3, S4 and Table S1). *CDKN2A* encodes tumour suppressors p16 and p14ARF and loss of this gene has been linked to resistance to targeted drugs (Mullighan et al., 2008). Analysis of single nucleotide variants (SNVs) revealed a small cluster of mutations clearly enriched in the trametinib resistant lines compared to POT (Figure 3D and S5). These mutations were also enriched in the gefitinib resistant lines, although to a lesser extent, potentially indicating a pre-existing subclone that is doubly-resistant to gefitinib and trametinib, although more strongly selected by trametinib. We speculate that these mutations are likely passenger hitchhikers of the *CDKN2A*-loss subclone.

**Figure 3:**
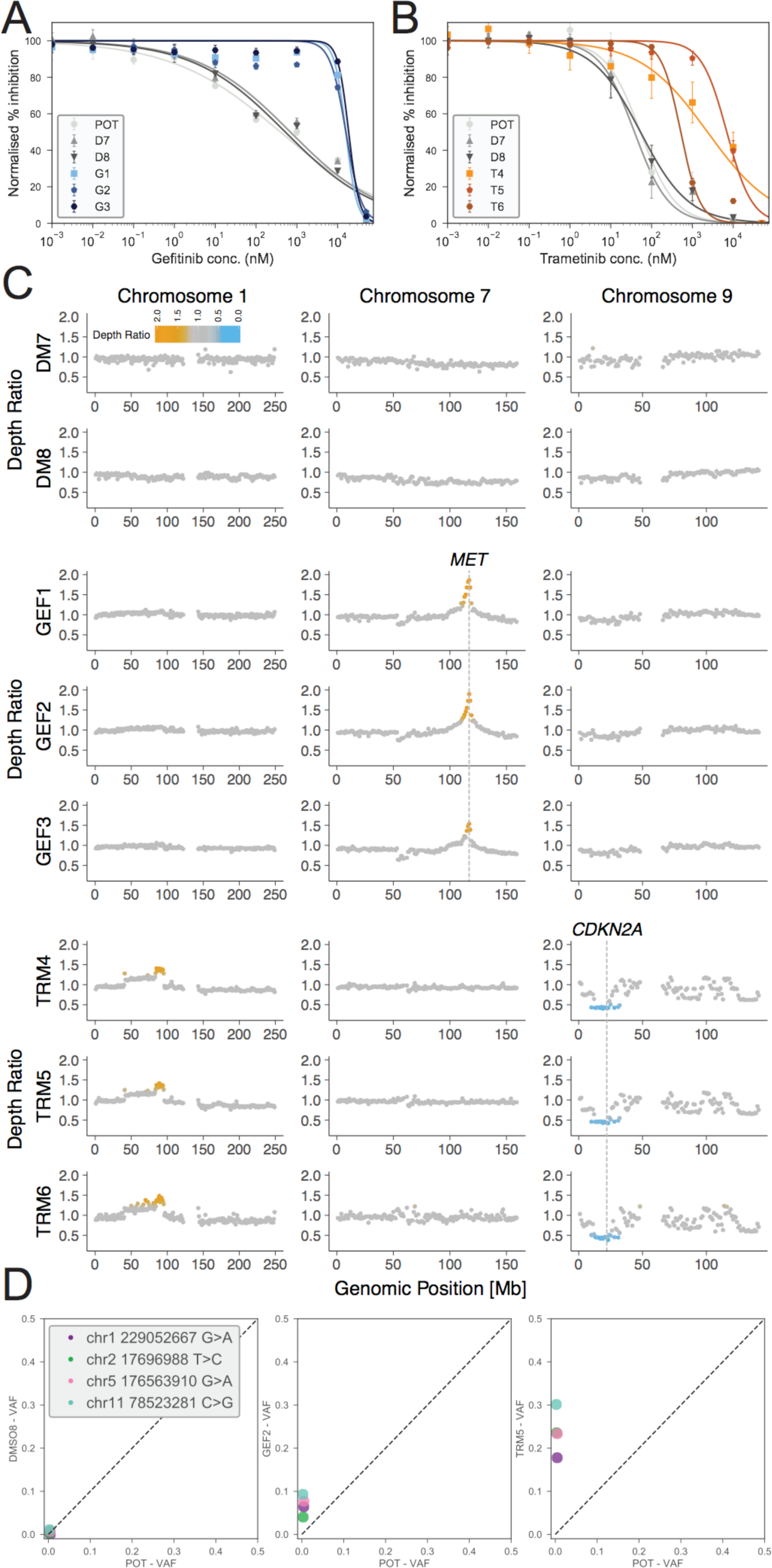
Characterisation of the resistant lines. **(A)** Dose-response curves of gefitinib resistant lines and **(B)** trametinib resistant lines versus DMSO demonstrate acquired resistance. **(C)** Relative copy number profiling of resistant lines compared to POT highlight MET amplification in GEF lines and 1p gain, 9p loss (including CDKN2A loss) in TRM lines. **(D)** Single-nucleotide variant analysis shows enrichment of a subclone containing 4 variants in TRM and partial enrichment in GEF of the same clone.

The fact that genomic alterations were consistent between evolved replicas but different for the two drugs suggest that multiple resistant subclones were already present in the initial population. Differential evolution and competition of these subclones under the two drugs also suggest a target for herding.

### Tracking clonal evolution in real time non-destructively

We next sought to more precisely quantify the temporal evolutionary dynamics. We profiled the barcodes of all samples using ultra-deep sequencing (Material and Methods). In comparison to the 2295 unique barcodes identified in the POT population, we found an average of 872 unique barcodes in the gefitinib treated lines and an average of 199 unique barcodes in the trametinib lines (Figure S6), indicating that drug exposure induced a strong selective bottleneck. We note that because of the single-cell barcoding, we expect multiple barcodes corresponding to each pre-existing subclone (i.e. multiple cells in the subclone have been barcoded with different barcodes). We considered a barcode as positively selected in a given replicate when its estimated growth rate was positive with respect to DMSO (Materials and Methods). We grouped barcodes with similar growth dynamics into ‘functional subclones’. We define pre-existing functional subclones as those having similar growth dynamics in more than one replica (Figure 4A and Material and Methods for details). Notably, we cannot exclude that each functional subclone may be composed of multiple genetically distinct subclones. This is not critical for our analysis as we are interested in drug response phenotypes, rather than individual genotypes.

**Figure 4:**
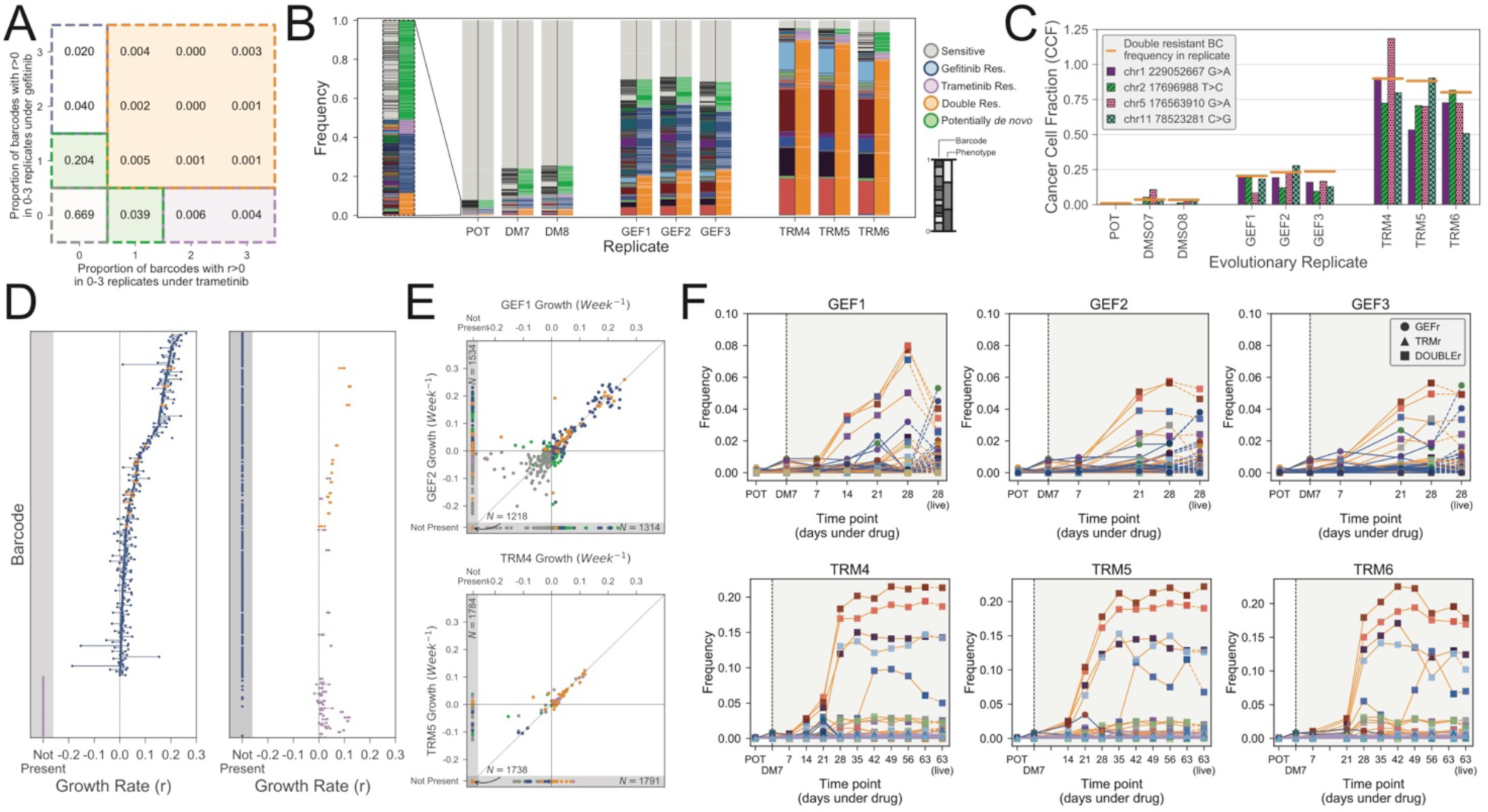
Barcodes reveal evolutionary dynamics over time. **(A)** Map of barcodes with drug response phenotype to define functional subclones. Values indicate the proportion of unique barcodes with positive growth rates across the given number of replicates of gefitnib exposure and trametinib exposure. **(B)** Barcode frequency distributions in each sample. Left hand bars show the frequency of each unique barcode. Barcode colours and ordering are identical between replicates. Right hand bars indicate the phenotypes assigned to each barcode. Phenotypes are determined by the set of evolutionary replicates in which a barcode exhibits a positive growth rate (Materials and Methods). Barcode and phenotype distributions are highly conserved between replicates, indicating repeatable evolution. **(C)** Cancer cell fraction estimates for the cluster of four SNVs identified form exome sequencing match barcode frequencies for the doubly-resistant phenotypes. **(D)** Growth rates for each barcode assigned to the GEF, TRM or double resistant phenotypes are shown under both the gefitinib (GEF1-3) and trametinib (TRM4-6) exposure. Points indicate the growth rates in the three replicates and lines connect these points to highlight variance. **(E)** Representative scatter plots show the concordance in barcode growth rates between evolutionary replicates. Points are coloured according to barcode phenotype, as in (B). **(F)** Temporal frequencies for the floating barcodes in each evolutionary replicate. Lines are coloured by phenotypes and marker colours correspond to the unique barcode as in (A). POT and DM7 measurement are harvested (live) populations as is the final time point, all others are floating barcode measurements. The temporal frequency dynamics are conserved both between time points within each replicate and between replicates. Moreover, the final samples (harvested population) largely match the last floating cells samples.

We identified five functional subclones with different growth dynamics (Figure 4B). The first group (grey) was the largest (87.2%) and represented largely clones that died under both drugs (sensitive) as well as clones for which the growth rate could not be determined because not found in the DMSO (Figure S7). The second group (blue) was resistant to gefitinib but sensitive to trametinib. The third group (purple) was resistant to trametinib but sensitive to gefitinib. The fourth group (orange) was doubly resistant to both drugs. Finally, the fifth group (green) was composed by a set of barcodes that were found in only one replica either of trametinib or gefitinib. This set could correspond to possible *de novo* resistant lineages. As this group comprises of barcodes all at low frequency, we focused on the majority of pre-existing resistant subclones that are relevant to evolutionary herding. We examined the frequency of barcodes and associated phenotypes in the POT versus the evolved lines. Strikingly, the frequencies of barcodes between replicas of a drug were highly similar, confirming that the initial conditions are a strong determinant of evolution under exposures to high drug concentrations (Figure 4B). Importantly, these results indicate that in this system, dynamics are deterministic and hence predictable.

We reasoned that the doubly resistant (orange) subclone could be the same carrying the SNVs found highly enriched in TRM and partially enriched in GEF using the exome sequencing analysis. We contrasted the barcodes frequency of the orange subclone with the SNV Cancer Cell Fraction (CCF) in each sample and found that these two independent measurements matched in all samples, including the POT, thus describing concordant evolutionary dynamics and suggesting that the SNVs and barcodes are in the same cells (Figure 4C).

Using mathematical modelling, we measured the growth rates of each barcode under each condition (see Material and Methods). This analysis confirmed that gefitinib resistant population was polyclonal, with a large *MET* amplified subclone (blue barcode group) composing ∼32.8% (average) of the population in GEF1-GEF3 and a relatively large initial population (∼2.4%) in the POT – see Figure 4B. This subclone was characterised by many barcodes with a positive growth rate under gefitinib but a negative growth rate under trametinib (Figure 4D – blue barcodes). We also found enrichment the multidrug resistant subclone (orange barcodes) that exhibited a positive growth rate under both gefitinib and trametinib. This subclone was found at mean frequency 22.4% in the GEF lines and 86.1% frequency in the TRM lines (Figure 4D – orange barcodes). This clone was smaller than the blue clone in the original POT population (∼0.91%) and therefore carried many fewer barcodes. There was also a small set of barcodes that were only enriched in the trametinib lines (Figure 4D – purple barcodes, ∼4.2% average in TRM lines, ∼0.57% in POT). The combined frequency of all enriched (resistant) barcodes in the initial population was 3.9%. The growth rates across replicates were highly similar (Figure 4E and S8). Hence, the barcode analysis supports the presence of pre-existing polyclonal drug resistance.

As part of our experimental design, we never re-plated cells following drug exposure in order to avoid strong stochastic drift effects due to sampling bias. As such, we could not take aliquots of cells for analysis throughout the experiment. To track evolution through time in a non-destructive way, we leveraged the large volume of media (560ml) that is changed every week. HCC827 is an adherent cell line, with cells that detach from the plate surface upon death. We collected pellets consisting of cells that had died within the week and extracted barcodes from each time point. We confirmed that pellets from supernatant collection were apoptotic/necrotic cells (Figure S9). Time-course barcodes allowed us to track the evolution under drug exposure without perturbing the system and at a resolution that is unparalleled (Figure 4F). Strikingly, this barcode analysis clearly shows an expansion of the subclones we identified in our analysis of the final populations, with the final time point of barcodes derived from supernatant cells being very similar to the final harvested populations (Figure 4F, line colours indicate phenotype, point colours indicate unique barcodes). This result seems partially counterintuitive, as one might expect the barcodes harvested from the dead cells not to correspond to a resistant clone. However, this phenomenon can be understood by consideration of the underlying evolutionary dynamics. At first, many barcodes are driven to extinction, because the majority of cells in the initial population are sensitive to the drug. At this stage, those cells present in the harvested media correspond to the thousands of different barcodes of sensitive cells (grey), none of which is common in the initial population, hence no enrichment is detected. As the resistant population grows, the contribution to the floating media becomes a mixture of sensitive cells being driven to extinction, and resistant cells turning over. At the end of the experiment, it is these resistant cells that are common and correspond to the few barcodes that are enriched. The frequencies of the clones stabilized after approximately 3 weeks of gefitnib exposure, and 6 weeks of trematinib exposure (Figure 4F). By comparing the time series barcode dynamics between replicates, we again see that the evolutionary dynamics are strikingly conserved, suggesting that the resistance dynamics are highly predictable (Figure 4F).

### Evolutionary trade-offs and collateral drug sensitivity

We have demonstrated quantitatively how gefitinib and trametinib herd the tumour population by expanding pre-existing resistant subclones. We sought to quantify the effect of evolutionary herding on the efficacy of second line drug treatment. Our genomic analysis predicts the existence of a MET amplified clone in the gefitinib treated lines and a separate CDKN2A-loss clone in the trametinib treated lines. We performed single-cell RNA sequencing on the POT sample, one gefitinib-treated replicate (GEF1) and one trametinib-treated replicate (TRM4). tSNE analysis confirmed that cells deriving from the same evolutionary replicates clustered closely together (Figure 5A). Colouring cells by the expression of these genes confirmed these predictions at the RNA level, including the subclonality of the MET amplification in the gefitinib resistant line (Figure 5B). Overall, the single-cell data provides a picture that confirms the clonal composition reported by the barcodes. Phosphoproteomic results validated our findings in terms of the functional effects of the drugs on the signalling pathways (Figure S11, see Material and Methods).

**Figure 5.**
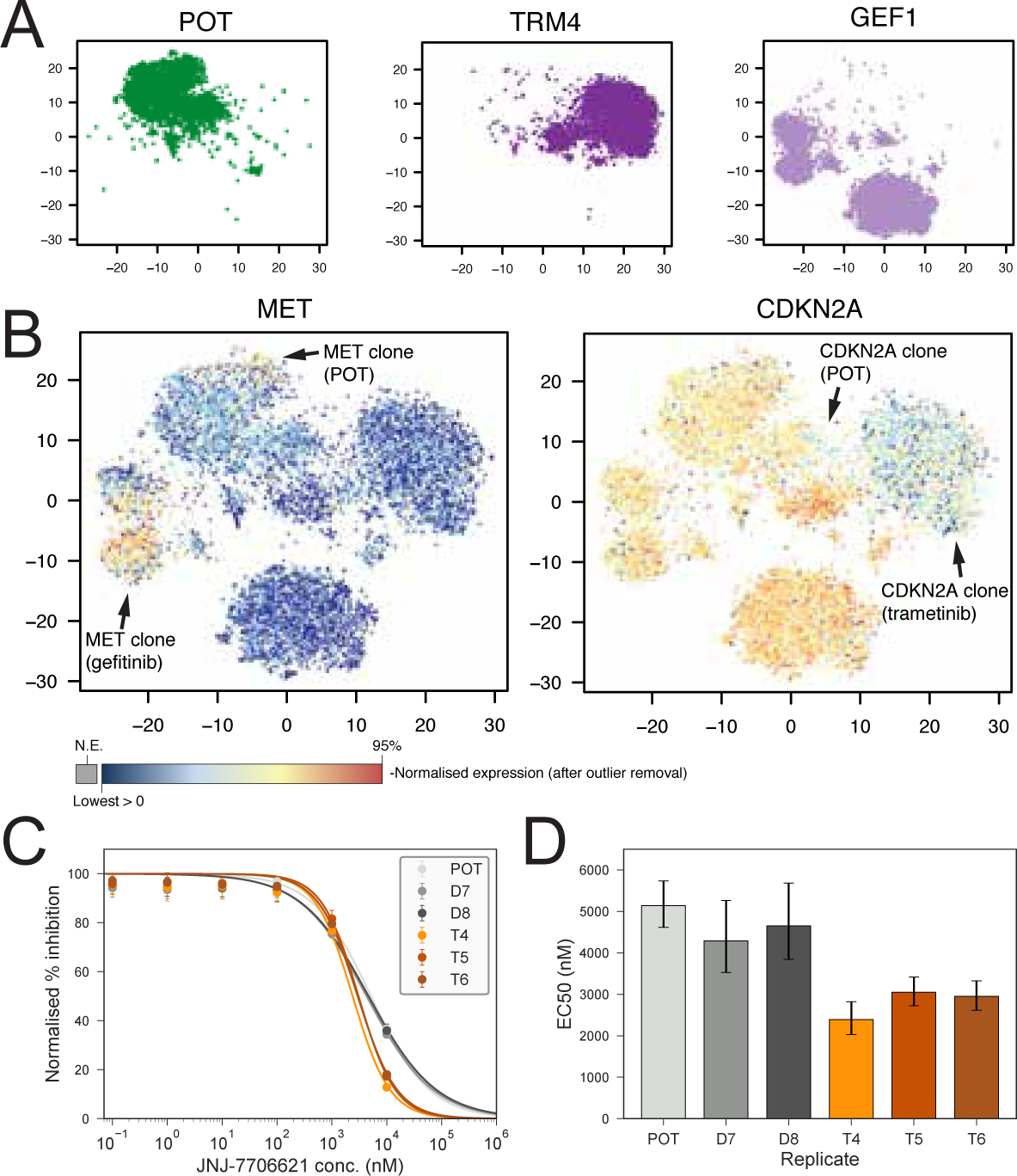
Herding leads to collateral drug sensitivity. **(A)** tSNE of single cell RNA sequencing coloured by replicate. **(B)** tSNE of single cell RNA sequencing coloured by decile of expression of MET and CDKN2A, different drug-resistant clones are indicated with arrows. **(C)** Dose response curves for the pan-CDK inhibitor JNJ-7706621 applied to the TRM lines (left). **(D)** EC50 concentrations for baseline versus evolved lines. Error bars denote 95% confidence intervals.

In light of these results, we reasoned that GEF the lines may exhibit collateral sensitivity to MET inhibition owing to the presence of a MET amplified subclone. As CDKN2A loss leads to upregulation of CDK2/4, we reasoned that inhibition of CDKs could prove effective in the TRM lines. Indeed, we found collateral sensitivity of TRM lines to JNJ-7706621, a pan-CDK inhibitor (Figure 5C), leading to half of the EC50 dose for trametinib evolved lines with respect to DMSO or POT (Figure 5D).

We then leveraged high throughput drug screening technology to assay sensitivity to a panel 485 compounds at each of four concentrations (20nM, 80nM, 200nM and 800nM). This screen revealed a total of 8 candidate collaterally sensitive drugs (Figure S10A,B) of which 6 were chosen for validation, including a cMET inhibitor BMS-777607 (see Material and Methods). However, none of these candidates exhibited collateral sensitivity upon validation (Figure S10C), suggesting that standard high throughput screening methodologies, developed for first line target identification, may not be ideal to study changes in drug efficacy following evolutionary herding.

Taken together, these results demonstrate the potential of evolutionary herding in which a first drug can change the clonal composition of the population, inducing stable collateral drug sensitivity. However, although evolutionary trade-offs can be exploited to obtain collateral drug sensitivities, these are often rare, and likely specific to the signalling pathways altered by the agent. This may explain why collateral sensitivities have proved difficult to identify, and supports the need for model systems that are better able to identify evolutionary trade-offs using tumour evolution and biologically informed drug screening.

## Discussion

The vast majority of metastatic cancers remain largely incurable. Treatment with standard approaches may extend survival (Zhang et al., 2009), but ultimately fails due to the emergence of resistant cells (McGranahan and Swanton, 2015). This is the natural consequence of a process of clonal evolution fuelled by intra-tumour heterogeneity (Greaves and Maley, 2012). Combining different drugs together at the same time has been investigated, but typically only improves survival by a few months, if any (Carrick et al., 2009; Delbaldo et al., 2004), and the narrow therapeutic window of cancer drugs leads to high toxicity in combinations, limiting the practicality of this approach. Instead, controlling the disease, rather than attempting to cure it, may be the only viable option in advanced cancers (Gatenby et al., 2009). Although this sounds radical in Oncology, disease management is well established in fields such as HIV (Ghosn et al., 2018) and antibiotic resistance (Nichol et al., 2015), as well as pest control (Alto et al., 2013; Neve et al., 2009; Oliveira et al., 2007). In cancer, different groups have explored this concept of ‘adaptive therapy’ (Gatenby et al., 2009) where drug dose is modulated in response to the underling evolutionary dynamics (Enriquez-Navas et al., 2016; Gallaher et al., 2018), with encouraging preliminary results in clinical trials (Zhang et al., 2017). Many adaptive approaches are based on ‘buffer therapy’, which exploits the fact that resistance often comes at a proliferative cost and hence resistant subpopulations may be at disadvantage in a drug-free environment (Hughes and Andersson, 2015b). This has been observed prospectively in colorectal cancer patients under EGFR inhibition, where KRAS-driven resistance seems to imply a cost, and KRAS subclones decrease in relative frequency if the drug is suspended (Siravegna et al., 2015). We have also observed this in our evolved lines treated with trametinib, which show significantly slower growth with respect to baseline. When resistance comes at a cost in a drug-free environment, the drug sensitive subpopulations can be used to “keep in check” drug resistant cells (Gatenby et al., 2009). This would explain the low prevalence in the POT of the CDKN2A-loss clone. Moreover, evolutionary game theory has been proposed as a conceptual framework for adaptive therapy (Staňková et al., 2018). In this study we evaluated an additional level of complexity, where we use more than one drug to study evolutionary trade-offs and costs of resistance not just in the original drug-free environment, but also under the pressure of secondary drugs. The specific type of adaptive therapy we investigated is evolutionary herding, which exploits the differences in fitness landscapes imposed by different drugs to control or prevent drug resistance.

Despite the conceptual elegance and promises of adaptive therapy however, current strategies are often based on *ad hoc* rules of thumb. The lack of reliable experimental model systems that recapitulate patient heterogeneity and clonal evolution is a major barrier for bringing adaptive therapies to the clinic. Here we presented a new approach for clonal herding where evolution can be tightly controlled, monitored and altered using drugs. This has the potential of paving the way to multidrug adaptive treatments.

Although we have attempted to design a model system that specifically aims at recapitulating the evolutionary dynamics of treatment resistance occurring in patients, our study has limitations. First, we do acknowledge that established cell lines may not recapitulate the dynamics of evolutionary herding in the patient. Second, we have used high concentrations of drugs that may not be always achievable in patients. Therefore, future studies will be needed that incorporate tumour microenvironment factors such as cancer-associated stromal and immune cells as well as different doses of drugs. Moreover, additional validation experiments will be needed prior to incorporating such models into clinical trial design.

Despite these limitations, model systems that recapitulate the temporal dynamics of human cancer evolution will shed new light on how to control drug resistance in advanced cancers, and open for the opportunity of personalised adaptive drug schedules that may achieve long-term control in advanced human cancers.

## Materials and Methods

### Cell line culture in HYPERflasks

HCC827 cell line was cultured in RPMI-1640 medium (Sigma-Aldrich) supplemented with 10% FBS (Sigma-Aldrich), 4 mM L-Glutamine (Sigma-Aldrich), 1% non-essential amino acids (Sigma-Aldrich), and 1% Penicillin-Streptomycin (Sigma-Aldrich). Cell line was confirmed to be *Mycoplasma* free using PCR-based method. Cell line was grown and expanded in High Yield PERformance Flasks (HYPERflask®) cell culture vessel (Corning). Medium was changed once a week and cells were harvested upon reaching ∼85% confluence.

### Barcoding of cell lines

The ClonTracer lentiviral barcode library construction and the generation of the lentivirus were previously described (*38*). The ClonTracer library a gift from Frank Stegmeier (Addgene #67267). HCC827 cell lines were cultured in normal growth media and barcoded by lentiviral infection using 0.8 μg/ml polybrene. For the majority of single cells to be infected with a single barcode a multiplicity of infection (MOI) of 0.1 corresponding to 10% infection was chosen, following lentiviral titration results. Following infection, 2,5 μg/ml puromycin was used for selection of cells infected with a barcode. Statistical analysis of the barcoding process suggests that <1% of cells were doubly barcoded and <0.1% of unique barcodes were received by multiple cells (Supplementary Methods).

### Generation of gefitinib and trametinib resistant cell lines

The 1 million barcoded HCC827 cells were expanded to approximately 120 million cells, harvested and frozen. Of these frozen cells, 4 million cells were thawed and again expanded to approximately 120 million cells. These cells were seeded into 8 HYPERflasks equally. Two HYPERflasks were grown under <0.0001% of DMSO for 6 days upon which they reached ∼85% confluence and were harvested as controls. The remaining 6 HYPERflasks were grown under normal growth media for one week upon which they were exposed to GI99 concentrations of gefitinib (Selleckchem) and trametinib (Selleckchem) (3 replicate flasks for each), for 4 and 9 weeks respectively. During this time, the medium and inhibitor were replenished weekly. The GI99 concentrations for gefitinib and trametinib were previously determined to be 40 nM and 100 nM (Figure S1). Cell counts were determined via the Countess II Automatic Cell Counter (ThermoFisher).

### Barcode amplification and next generation library preparation

Barcoded HCC827 cell lines and human PDOs were harvested and pelleted. Genomic DNA isolation was performed using DNeasy Blood and Tissue DNA extraction kit (Qiagen) according to manufacturer’s recommendations. Half of the conditioned media from each HYPERflasks were centrifuged at 1,700 x *g* and pelleted. Quantification of genomic DNA was carried out using Qubit (Life Technologies). Amplicon PCR reaction was performed using 2x Accuzyme mix (Bioline) and 20 ng of DNA to amplify the barcode using the previously published primers sequences (Bhang et al., 2018):

Forward: ACTGACTGCAGTCTGAGTCTGACAG

Reverse: CTAGCATAGAGTGCGTAGCTCTGCT

Following detection of 80-bp PCR product including the 30-bp semi-random barcode and after purification, NGS libraries were prepared using the NEBnext Ultra II DNA library preparation kit for Ilumina (New England Biolabs) according to manufacturer’s recommendations. Libraries were quantified using Qubit (Life Technologies) and KAPA library quantification kit (KAPA Biosystems), as well as TapeStation (Agilent Genomics). Library preparation was not successful for DNA extracted at four floating cell time points (GEF2-F2, GEF3-F2, TRM5-F1 and TRM6-F3). NGS was performed in house using MiSeq (Ilumina).

### Barcode bioinformatics analysis

FastQ files were first filtered to extract those reads with quality score >20 in all positions. Reads matching potential barcodes were extracted from FastQ files by use of a regular expression matching 12 bases of the forward barcode primer, followed by 30 base pairs, followed by 12 bases of the reverse barcode primer. To account for potential errors arising from PCR amplification or mutation, similar barcodes were merged via a novel method (outlined in the Supplementary Methods) that assigns each barcode to a representative matching the known weak/strong base pair pattern by consideration of the Hamming distance between barcodes.

To assign a phenotype to each barcode we first determined an approximate growth rate under each condition by consideration of the frequencies. We assumed that the frequency of each barcode in the DMSO replicates was representative of the frequency in the drug-treated replicated GEF1-GEF3, TRM4-TRM6 prior to the introduction of drug. To ensure a conservative estimate of the growth rate, we estimated the initial frequency as 𝑓_0_ = *Max*(𝑓*_D7_*, 𝑓*_D8_*) where 𝑓*_D7_*, 𝑓*_D8_* denote the frequency of the barcode in the lines DMSO7 and DMSO8 respectively. Denote the barcode frequency in a given replicate following drug exposure, expansion and harvesting by 𝑓_*R*_. We estimated the growth rate of the barcode under drug exposure as

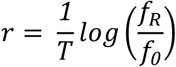

where *log* denotes the natural logarithm and *T* denotes the time between drug exposure and harvesting the cells (*T* = *4weeks* for gefitinib, *T* = *9weeks* for trametinib).

Phenotypes were then assigned according to the number of gefitinib and trametinib evolutionary replicates in which the barcode exhibited a positive growth rate. As a barcode can appear extinct in a given replicate either because it has negative growth rate, because the specific barcode was never seeded to that replicate, or because of drift, we determined barcode phenotypes as follows. Where a barcode exhibited positive growth rate in both 1+ GEF and 1+ TRM replicates, the barcode was designated as double-resistant. Where a barcode exhibited positive growth rate in 2+ GEF lines but no TRM lines it is designated gefitinib resistant / trametinib sensitive. Likewise, where a barcode exhibited positive growth rate in 2+ TRM lines but no GEF lines it is designated trametinib resistant / gefitinib sensitive. Where a barcode exhibits positive growth rate in a single replicate (GEF or TRM) it is designated as *putatively de novo* resistance. Other barcodes with measured growth rate are designated sensitive. Finally, some barcodes are designated as having undetermined phenotype where a barcode is not detected in DMSO7 or DMSO8 (potentially due to loss at seeding) but observed in a replicate, as a growth rate cannot be determined. Figure 4A shows a schematic of the phenotype mapping along with the proportion of unique barcodes assigned to each phenotype. Moreover, we compared a previous ‘POT’ baseline sample with the POT used in this experiment, after it has been frozen, stored and then thawed. Figure S12 shows that barcodes are highly consistent in terms of proportion in the two samples, with a proportion of barcodes that are always missed by sequencing, which implies a binomial sampling of the barcodes.

### Whole Exome Sequencing

Nine whole exome sequencing libraries were prepared from 200ng of genomic DNA using the Agilent SureSelect HT2 Human All Exon_V6 kit following the manufacturer’s instructions. The libraries were pooled and sequenced on the Illumina NovaSeq platform. The median (of medians) coverage achieved was 161x (min 43x, max 218x), see Table S2.

#### Mutation Calling

Trimming was performed with Skewer v0.1.126. Reads with mean a quality value greater than 10 prior to trimming and a minimum read length of 35 following trimming were kept. All others were discarded. Trimmed reads were aligned to the full human reference genome hg19 with the Burrows-Wheeler Aligner tool (bwa-mem, v0.7.15). PCR duplicates were marked using Picard tools (v2.8.1). Mutations were jointly called for all samples together using Platypus v0.8.1(Rimmer et al., 2014). The extent of selection was determined by identifying SNVs exhibiting a 10x enrichment in VAF in the treated lines (GEF1-GEF3, TRM4-TRM6) over the POT line. This analysis yielded a cluster of four SNVs exhibiting enrichment corroborating that predicted by the barcode enrichment analysis.

##### Copy Number Analysis

Heterozygous single nucleotide polymorphisms (SNPs) in the exome sequencing of the cell lines were identified using allelecount v3.0.1 (www.github.com/cancerit/alleleCount). Here we counted bases at SNP locations that have a global minor allele frequency between 0.1 and 0.2 (min. genomic position 100,000 bp) in dbSNP build 132 (Sherry et al., 2001) and overlap with the target regions of the exome panel for all autosomes. We calculate B-allele frequency (BAF) by dividing the highest base pair count by the total coverage at the SNP loci. These values were randomly subtracted from 1 to simulate the random assignment of the A and B allele. Log R ratio (LRR) was calculated as the log base 2 of the coverage of each SNP loci normalised by subtracting the global median LRR value.

To identify segments of copy number alterations (CNAs) we smoothed and segmented the LRRs of each sample using DNAcopy. In order to calculate the mean heterozygous major allele frequency in each segment we required a test for distinguishing between segments with pure loss-of-heterozygosity (LOH) and segments containing heterozygous SNPs. We identified segments with heterozygosity by counting the numbers of SNPs in each segment with a major allele frequency less than 0.9. We then performed an exact binomial test in which the alternative hypothesis was that more than 5% of the segment contains heterozygous SNPs (p < 0.05). For those segments in which the null hypothesis was rejected, the median heterozygous major allele frequency value was used to represent the allelic (im)balance of the segment.

Using the ASCAT equations (Van Loo et al., 2010), we assumed each sample was pure (rho = 1) and solved the ploidy of each sample (psi) by calculating the distance of the continuous major and minor copy number values of all segments from their nearest integer states across a range of psi values that are realistic for tumour ploidy (1.5 to 5.5). The psi value that produced the smallest distance from integers in all segments was taken as the ploidy solution. This was ∼3 for all cell lines as the cell line HCC827 is known to be triploid (Engelman et al., 2007).

We additionally calculated GC content normalised depth ratios between each treated cell line and the parental population (POT) using Sequenza (Favero et al., 2014). To calculate segments of differential copy number status, we subset the loci by their global minor allele frequency in dbSNP build 132 and segmented the depth ratios using DNAcopy as described previously.

### Digital Droplet PCR

Genomic DNA isolation for ddPCR was performed using DNeasy Blood and Tissue DNA extraction kit (Qiagen) according to manufacturer’s recommendations. Quantification of gDNA was carried out using Qubit (Life Technologies). DdPCR was performed on a QX200 ddPCR machine (Bio-Rad). Copy number assay was performed using 3 ng gDNA as a template and commercially available probes for *MET* (dHSACP2500321, FAM, Bio-Rad) and *NSUN3* (dHSACP2506682, HEX, Bio-Rad) as a reference gene. PCR reactions were performed using 3ng of DNA, 10 μl of 2xSupermix in a total volume of 20 μl. Automated droplet generator (Bio-Rad) was used to generate ∼20,000 droplets for partition of PCR reactions. Negative controls with no DNA and positive control DNA extracted from a cell line with previously reported CN were included. QuantaSoft v1.3.2.0 software was used for *MET* CN analysis. Copy number status of *NSUN3* was assumed to be 3 (triploid) and this was confirmed by copy number analysis in exome sequencing data.

### High-throughput drug screening

Cells from POT, DMSO7, DMSO8, GEF1-3 and TRM4-6 were tyripsined and counted. 1,000 cells per well were seeded in 384-well plates (Corning). Cells were grown in a 37°C and 5% CO_2_ incubator overnight. A panel of 485 agents (Table S3) was prepared in 4 different concentrations (20 nM, 80 nM, 200 nM and 800 nM) and dispensed per well using Echo 555 liquid handler (Labcyte Inc.). After 3 days of treatment with agents, cells were incubated with 10% CellTiter-Blue cell viability reagent (Promega) for 4 hours in a 37°C and 5% CO_2_ cell culture incubator. Finally, EnVision (PerkinElmer) plate reader was used to obtain readings.

Hit identification was performed separately for each of the four drug concentrations. First, for each replicate, normalised percentage inhibitions (PCI) for each compound were derived using the average fluorescence of 14 empty wells as a negative control and the average fluorescence of 14 wells seeded with cells but no drug as a positive control. To improve statistical power, we grouped the PCIs for the two POT controls and two DMSO lines together as a control data set comprising four data points per compound, per concentration. Likewise, the data for the three evolutionary replicates under each drug (GEF1-3 and TRM4-6) were grouped. Potential collaterally sensitive second line therapies were identified as those exhibiting a significant change in mean PCI (as determined by a two-tailed t-test on the PCIs at an 〈=0.05 significance threshold) that was greater than a 5%. To account for spurious hits arising from multiple hypothesis testing we considered only compounds resulting in hits at two or more concentrations. This analysis yielded eight candidate therapies for validation (Figure S8), namely Teniposide, Mycophenolate Mofetil, Fludarabine, BMS-777607, Tosedostat (CHR2797), Panobinostat (LBH589), ENMD-2076, PF-03814735. Of these we opted to validate six, excluding Mycophenolate Mofetil and Fludarabine. The six drugs for validation were purchased from Selleckchem.

### High-throughput drug screen validation

POT, 2 DMSO, gefitinib and trametinib resistant cell lines were trypsinised and counted. Between 500 and 10,000 cells per well were seeded in 96-well standard plates (Corning). Following overnight incubation in a 37°C and 5% CO_2_ cell culture incubator, average 10-fold changing dose of 10 concentrations from each inhibitor were used. 3 days post inhibitor treatment for all of the drugs validated, with an exception of 10 days for trametinib, 10% CellTiter-Blue cell viability reagent (Promega) was applied. After overnight of incubation with 10% CellTiter-Blue in a 37°C and 5% CO_2_ cell culture incubator, readings were obtained using EnVision (PerkinElmer) plate reader.

To derive dose-response curves, normalised percentage growth was derived from OD readings by normalisation to six positive control (drug-free growth) and six negative control (empty) wells. A two parameter (ec50, hill coefficient) log-logistic dose response curve was then fitted to the data via non-linear least squares regression.

### Luminex phospohoprotein Assay

POT, DMSO7, GEF1 and TRM4 cell lines were tyripsinied and counted. Following seeding of 300, 000 cells per well in 6-well plates and incubation in a 37°C and 5% CO_2_ cell culture incubator overnight, 3 biological replicates of each cell lines were treated with DMSO, 40nM of gefitinib and 100nM of trametinib for 1 hour. After the incubation under those conditions, cells were tyripsinised and centrifuged at 1,500 rpm to generate cell pellets. Cell pellets were lysed using MDS Tris Lysis Buffer (Meso Scale Diagnostics) containing phosphatase inhibitor I (Sigma-Aldrich), phosphatase inhibitor II (Sigma-Aldrich), protease inhibitor (Cell Signalling Technology). Protein content of lysed samples was quantified using BCA assay (Sigma-Aldrich). MILLIPLEX MAP Akt/mTOR phosphoprotein kit, MILLIPLEX MAPK/SAPK signalling kit, MILLIPLEX MAP RTK phosphoprotein kit (48-611MAG, 48-660MAG, HPRTKMAG-01K respectively, MerckMillipore) were combined with the following singleplex magnetic bead sets to produce three multiplex Luminex assays; Total HSP27, GAPDH (46-702MAG, 46-710MAG, 46-623MAG, 46-641MAG, 46-608MAG, 46-667Mag, MerckMilipore). Bio-Plex Pro phosphor-PDGFRb and Akt (Thr308) (171-V50018M, 171-V50002, Bio-Rad) were combined into a triplex assay. Manufacturer’s recommendations were followed. Phosphoprotein levels were measured on the Luminex 200 system utilizing xPOTENT c3.1 software.

EGFR phosphorilation was highly downregulated under gefitinib and even in the absence of drug in GEF evolved lines, suggesting a stable phenotype where EGFR signalling has been lost due to clonal evolution (Figure S10A). MET phisphorilation was upregulated only in MET amplified GEF lines, as expected (Figure S10B). MEK phorphorilation was variable (Figure S10C), however we confirmed ERK/MAPK downregulation under trametinib (Figure S10D), a strong indicator that the drug is inhibiting the MEK pathway.

### Floating barcodes harvesting

To track evolution through time, we leveraged the large volume of media (560ml) that must be changed each week to maintain the HYPERflask culture system. HCC827 is an adherent cell line, with cells that detach from the plate surface upon death. By spinning the spent media in a centrifuge at 12,000 rpm for 10 minutes, we collected pellets consisting of cells that had died within the week. We extracted barcodes from these intermediate time points for each of the gefitinib exposed lines (weekly for 4 weeks) and for each of the trametinib resistant exposed lines (weekly for 9 weeks). These barcodes permitted us to track the evolution of each cell lineage, under each drug exposure, without the need for re-plating, and with a temporal resolution that is unparalleled. Apoptotic barcoded cells were extracted using DNeasy Blood and Tissue DNA extraction kit (Qiagen).

### Single cells RNA profiling

#### Sample preparation

Single cells were prepared from POT, GEF1 and TRM4 cells. After centrifugation, single cells were washed with PBS and were re-suspended with a buffer (Ca^++^/Mg^++^ free PBS + 0.04% BSA) at 1,000 cells/µl.

#### Sequencing

Viability was confirmed to be > 90% in all samples using acridine orange/propidium iodide dye with LUNA-FL Dual Fluorescence Cell Counter (Logos Biosystems, L20001). Single cell suspensions were loaded on a Chromium Single Cell 3’ Chip (10X Genomics) and were run in the Chromium Controller to generate single-cell gel bead-in-emulsions using the 10X genomics 3’ Chromium v2.0 platform as per manufacturer’s instructions. Single-cell RNA-seq libraries were prepared according to the manufacturer’s protocol and the library quality was confirmed with a Bioanalyzer High-Sensitivity DNA Kit (Agilent, 5067-4627) and a Qubit dsDNA HS Assay Kit (ThermoFisher, Q32851). Samples were pooled up to three and sequenced on an Illumina HiSeq 4000 according to standard 10X Genomics protocol.

#### Data analysis

cellRanger (v2.1.1) was run on the raw data using GRCh38 annotation (v1.2.0). Output from cellRanger was loaded into the statistical computing environment R v3 (www.r-project.org) through the function *load_cellranger_matrix_h5* from package *cellranger* (v1.1.0; genome = “GRCh38”). Datasets were merged according to gene names. Before normalization, a series of filtering steps was performed. Only those cells showing at least 1,500 detected genes and 5,000 UMIs were considered for further analyses (Torre et al., 2018). Reads mapping on mitochondrial genes were excluded. After that, data were imported in *Seurat* (v2.3.4) (Butler et al., 2018) and scaled (*NormalizeData* function using normalization.method = “LogNormalize”, scale.factor = 10000, followed by the *ScaleData* function). A further filtering step was performed based on the cumulative level of expression (the sum of the Seurat-scaled values) of three housekeeping genes (GAPDH, RPL26 and RPL36) (Lin et al., 2018). Manual inspection of these values versus the number of UMIs per cell (or the number of genes with non-zero expression per cell) revealed no significant correlation between the two. Nevertheless, a number of cells showed extremely low expression of these genes, so those in the bottom 1% were excluded from further analyses. At last, genes expressed in less than 20 cells were also excluded. Linear normalization and scaling were performed again on the filtered, raw data. Variable genes were identified using the *FindVariableGenes* function of *Seurat* (mean.function = ExpMean, dispersion.function = LogVMR, x.low.cutoff = 0.01, x.high.cutoff = 6, y.cutoff = 0.01, num.bin = 100). Principal component analysis (PCA) was run using variable genes as input and, based on *p*-values estimated by the *JackStraw* function, the top 44 components were kept. These components were used as input for further dimensionality reduction (using t-Distributed Stochastic Neighbor Embedding; t-SNE) through the *RunTSNE* function (perplexity = 50, do.fast = TRUE, seed.use=44). Clusters were then identified using *FindClusters* (resolution = 0.6).

## Acknowledgments

A.S. is supported by the Wellcome Trust (202778/B/16/Z), Cancer Research UK (A22909) and the Chris Rokos Fellowship in Evolution and Cancer. We acknowledge funding from the National Institute of Health (NCI U54 CA217376) to A.S. and C.C.M. This work was also supported a Wellcome Trust award to the Centre for Evolution and Cancer (105104/Z/14/Z). U.B. is supported by The NIHR (RP-2016-07-28), CRUK (A25128, A22897) and by an NIHR Biomedical Research Centre Grant to the Institute of Cancer Research and The Royal Marsden NHS Foundation Trust. C.C.M. was supported in part by NIH grants U54 CA217376, P01 CA91955, R01 CA170595, R01 CA185138 and R01 CA140657 as well as CDMRP Breast Cancer Research Program Award BC132057 and an Arizona Investigator Grant ADHS18-198847. We thank Michael Hubank, Eleni Koutroumanidou and Debbie Hughes for technical support.

## Data Access

Sequence data have been deposited at the European Genome-phenome Archive (EGA), which is hosted by the EBI and the CRG, under accession number EGAS00001003200. Further information about EGA can be found on https://ega-archive.org.

## References

1. Alto, B.W., Lampman, R.L., Kesavaraju, B., and Muturi, E.J. (2013). Pesticide-Induced Release From Competition Among Competing Aedes aegypti and Aedes albopictus (Diptera: Culicidae). J Med Entomol 50, 1240–1249.

2. Bhang, H.-E.C., Ruddy, D.A., Krishnamurthy Radhakrishna, V., Caushi, J.X., Zhao, R., Hims, M.M., Singh, A.P., Kao, I., Rakiec, D., Shaw, P., et al. (2015). Studying clonal dynamics in response to cancer therapy using high-complexity barcoding. Nat. Med. –.

3. Butler, A., Hoffman, P., Smibert, P., Papalexi, E., and Satija, R. (2018). Integrating single-cell transcriptomic data across different conditions, technologies, and species. Nature Biotechnology 36, 411–420.

4. Carrick, S., Parker, S., Thornton, C.E., Ghersi, D., Simes, J., and Wilcken, N. (2009). Single agent versus combination chemotherapy for metastatic breast cancer. Cochrane Database of Systematic Reviews 34, 27.

5. Delbaldo, C., Michiels, S., Syz, N., Soria, J.-C., Le Chevalier, T., and Pignon, J.-P. (2004). Benefits of adding a drug to a single-agent or a 2-agent chemotherapy regimen in advanced non-small-cell lung cancer: a meta-analysis. - PubMed - NCBI. Jama 292, 470–484.

6. Diaz, L.A., Williams, R.T., Wu, J., Kinde, I., Hecht, J.R., Berlin, J., Allen, B., Bozic, I., Reiter, J.G., Nowak, M.A., et al. (2012). The molecular evolution of acquired resistance to targeted EGFR blockade in colorectal cancers. Nature 486, 537–540.

7. Domínguez-Vigil, I.G., Moreno-Martínez, A.K., Wang, J.Y., Roehrl, M.H.A., and Barrera-Saldaña, H.A. (2018). The dawn of the liquid biopsy in the fight against cancer. Oncotarget 9, 2912–2922.

8. Engelman, J.A., Zejnullahu, K., Mitsudomi, T., Song, Y., Hyland, C., Park, J.O., Lindeman, N., Gale, C.-M., Zhao, X., Christensen, J., et al. (2007). MET Amplification Leads to Gefitinib Resistance in Lung Cancer by Activating ERBB3 Signaling. Science 316, 1039–1043.

9. Enriquez-Navas, P.M., Kam, Y., Das, T., Hassan, S., Silva, A., Foroutan, P., Ruiz, E., Martinez, G., Minton, S., Gillies, R.J., et al. (2016). Exploiting evolutionary principles to prolong tumor control in preclinical models of breast cancer. Science Translational Medicine 8, 327ra24.

10. Favero, F., Joshi, T., Marquard, A.M., Birkbak, N.J., Krzystanek, M., Li, Q., Szallasi, Z., and Eklund, A.C. (2014). Sequenza: allele-specific copy number and mutation profiles from tumor sequencing data. Annals of Oncology 26, 64–70.

11. Fuentes-Hernandez, A., Plucain, J., Gori, F., Pena-Miller, R., Reding, C., Jansen, G., Schulenburg, H., Gudelj, I., and Beardmore, R. (2015). Using a sequential regimen to eliminate bacteria at sublethal antibiotic dosages. PLoS Biol. 13, e1002104.

12. Gallaher, J.A., Enriquez-Navas, P.M., Luddy, K.A., Gatenby, R.A., and Anderson, A.R.A. (2018). Spatial Heterogeneity and Evolutionary Dynamics Modulate Time to Recurrence in Continuous and Adaptive Cancer Therapies. Cancer Res. 78, 2127–2139.

13. Gatenby, R.A., Silva, A.S., Gillies, R.J., and Frieden, B.R. (2009). Adaptive Therapy. Cancer Res. 69, 4894–4903.

14. Ghosn, J., Taiwo, B., Seedat, S., Autran, B., and Katlama, C. (2018). HIV. The Lancet.

15. Gillies, R.J., Verduzco, D., and Gatenby, R.A. (2012). Evolutionary dynamics of carcinogenesis and why targeted therapy does not work. Nat. Rev. Cancer 12, 487–493.

16. Gottesman, M.M., and Pastan, I. (1993). Biochemistry of multidrug resistance mediated by the multidrug transporter. - PubMed - NCBI. Annu. Rev. Biochem. 62, 385–427.

17. Greaves, M., and Maley, C.C. (2012). Clonal evolution in cancer. Nature 481, 306–313.

18. Hall, M.D., Handley, M.D., and Gottesman, M.M. (2009). Is Resistance Useless? Multidrug Resistance and Collateral Sensitivity. Trends in Pharmacological Sciences 30, 546–556.

19. Hata, A.N., Niederst, M.J., Archibald, H.L., Gomez-Caraballo, M., Siddiqui, F.M., Mulvey, H.E., Maruvka, Y.E., Ji, F., Bhang, H.-E.C., Krishnamurthy Radhakrishna, V., et al. (2016). Tumor cells can follow distinct evolutionary paths to become resistant to epidermal growth factor receptor inhibition. - PubMed - NCBI. Nat. Med. 22, 262–269.

20. Holohan, C., Van Schaeybroeck, S., Longley, D.B., and Johnston, P.G. (2013). Cancer drug resistance: an evolving paradigm. Nat. Rev. Cancer 13, 714–726.

21. Hughes, D., and Andersson, D.I. (2015a). Evolutionary consequences of drug resistance: shared principles across diverse targets and organisms. Nat. Rev. Genet. 16, 459–471.

22. Hughes, D., and Andersson, D.I. (2015b). Evolutionary consequences of drug resistance: shared principles across diverse targets and organisms. Nat. Rev. Genet. 16, 459–471.

23. Imamovic, L., and Sommer, M.O.A. (2013). Use of Collateral Sensitivity Networks to Design Drug Cycling Protocols That Avoid Resistance Development. Science Translational Medicine 5, 204ra132–204ra132.

24. Kirkman, L.A., Zhan, W., Visone, J., Dziedziech, A., Singh, P.K., Fan, H., Tong, X., Bruzual, I., Hara, R., Kawasaki, M., et al. (2018). Antimalarial proteasome inhibitor reveals collateral sensitivity from intersubunit interactions and fitness cost of resistance. Proc. Natl. Acad. Sci. U.S.a. 115, 201806109–E201806870.

25. Lin, Y., Ghazanfar, S., Strbenac, D., Wang, A., Patrick, E., Lin, D., Speed, T., Yang, J., and Yang, P. (2018). Evaluating stably expressed genes in single cells. bioRxiv 229815.

26. Luria, S.E., and Delbrück, M. Mutations of bacteria from virus sensitivity to virus resistance. Genetics 28, 491–511.

27. McGranahan, N., and Swanton, C. (2015). Biological and therapeutic impact of intratumor heterogeneity in cancer evolution. Cancer Cell 27, 15–26.

28. McGranahan, N., and Swanton, C. (2017). Clonal Heterogeneity and Tumor Evolution: Past, Present, and the Future. Cell 168, 613–628.

29. Meacham, C.E., and Morrison, S.J. (2013). Tumour heterogeneity and cancer cell plasticity. Nature 501, 328–337.

30. Merlo, L.M.F., Pepper, J.W., Reid, B.J., and Maley, C.C. (2006). Cancer as an evolutionary and ecological process. Nat. Rev. Cancer 6, 924–935.

31. Misale, S., Yaeger, R., Hobor, S., Scala, E., Janakiraman, M., Liska, D., Valtorta, E., Schiavo, R., Buscarino, M., Siravegna, G., et al. (2012). Emergence of KRAS mutations and acquired resistance to anti-EGFR therapy in colorectal cancer. Nature 486, 532–536.

32. Mullighan, C.G., Williams, R.T., Downing, J.R., and Sherr, C.J. (2008). Failure of CDKN2A/B (INK4A/B-ARF)-mediated tumor suppression and resistance to targeted therapy in acute lymphoblastic leukemia induced by BCR-ABL. Genes Dev. 22, 1411–1415.

33. Neve, P., Vila-Aiub, M., and Roux, F. (2009). Evolutionary-thinking in agricultural weed management. New Phytologist 184, 783–793.

34. Nichol, D., Robertson-Tessi, M., Jeavons, P., and Anderson, A.R.A. (2016). Stochasticity in the Genotype-Phenotype Map: Implications for the Robustness and Persistence of Bet-Hedging. Genetics 204, 1523–1539.

35. Nichol, D., Jeavons, P., Fletcher, A.G., Bonomo, R.A., Maini, P.K., Paul, J.L., Gatenby, R.A., Anderson, A.R.A., and Scott, J.G. (2015). Steering Evolution with Sequential Therapy to Prevent the Emergence of Bacterial Antibiotic Resistance. PLoS Comput. Biol. 11, e1004493.

36. Nichol, D., Rutter, J., Bryant, C., Jeavons, P., Anderson, A., Bonomo, R., and Scott, J. (2017). Collateral sensitivity is contingent on the repeatability of evolution. bioRxiv 185892.

37. Oliveira, E.E., Guedes, R.N.C., Tótola, M.R., and De Marco, P., Jr. (2007). Competition between insecticide-susceptible and -resistant populations of the maize weevil, Sitophilus zeamais. - PubMed - NCBI. Chemosphere 69, 17–24.

38. Pao, W., Miller, V.A., Politi, K.A., Riely, G.J., Somwar, R., Zakowski, M.F., Kris, M.G., and Varmus, H. (2005). Acquired resistance of lung adenocarcinomas to gefitinib or erlotinib is associated with a second mutation in the EGFR kinase domain. - PubMed - NCBI. PLoS Med 2, e73.

39. Pál, C., Papp, B., and Lázár, V. (2015). Collateral sensitivity of antibiotic-resistant microbes. Trends in Microbiology 23, 401–407.

40. Pluchino, K.M., Hall, M.D., Goldsborough, A.S., Callaghan, R., and Gottesman, M.M. (2012). Collateral sensitivity as a strategy against cancer multidrug resistance. Drug Resist. Updat. 15, 98–105.

41. Rimmer, A., Phan, H., Mathieson, I., Iqbal, Z., Twigg, S.R.F., Wilkie, A.O.M., McVean, G., and Lunter, G. (2014). Integrating mapping-, assembly- and haplotype-based approaches for calling variants in clinical sequencing applications. Nature Genetics 46, 912–918.

42. Shaffer, S.M., Dunagin, M.C., Torborg, S.R., Torre, E.A., Emert, B., Krepler, C., Beqiri, M., Sproesser, K., Brafford, P.A., Xiao, M., et al. (2017). Rare cell variability and drug-induced reprogramming as a mode of cancer drug resistance. Nature 546, 431–435.

43. Sharma, S.V., Lee, D.Y., Li, B., Quinlan, M.P., Takahashi, F., Maheswaran, S., McDermott, U., Azizian, N., Zou, L., Fischbach, M.A., et al. (2010). A chromatin-mediated reversible drug-tolerant state in cancer cell subpopulations. - PubMed - NCBI. Cell 141, 69–80.

44. Sherry, S.T., Ward, M.H., Kholodov, M., Baker, J., Phan, L., Smigielski, E.M., and Sirotkin, K. (2001). dbSNP: the NCBI database of genetic variation. Nucleic Acids Res. 29, 308–311.

45. Shibue, T., and Weinberg, R.A. (2017). EMT, CSCs, and drug resistance: the mechanistic link and clinical implications. Nat Rev Clin Oncol 14, 611–629.

46. Siravegna, G., Mussolin, B., Buscarino, M., Corti, G., Cassingena, A., Crisafulli, G., Ponzetti, A., Cremolini, C., Amatu, A., Lauricella, C., et al. (2015). Clonal evolution and resistance to EGFR blockade in the blood of colorectal cancer patients. Nat. Med. 21, 795–801.

47. Staňková, K., Brown, J.S., Dalton, W.S., and Gatenby, R.A. (2018). Optimizing Cancer Treatment Using Game Theory: A Review. JAMA Oncol.

48. Torre, E., Dueck, H., Shaffer, S., Gospocic, J., Gupte, R., Bonasio, R., Kim, J., Murray, J., and Raj, A. (2018). Rare Cell Detection by Single-Cell RNA Sequencing as Guided by Single-Molecule RNA FISH. Cell Systems 6, 171–179.e175.

49. Van Loo, P., Nordgard, S.H., Lingjærde, O.C., Russnes, H.G., Rye, I.H., Sun, W., Weigman, V.J., Marynen, P., Zetterberg, A., Naume, B., et al. (2010). Allele-specific copy number analysis of tumors. Proc. Natl. Acad. Sci. U.S.a. 107, 16910–16915.

50. Wright, S. The roles of mutation, inbreeding, crossbreeding and selection in evolution. Proceedings of the Sixth International Congress of Genetics, Vol. 1 (1932), Pp. 356–366 1, 356–366.

51. Zhang, J., Yang, P.L., and Gray, N.S. (2009). Targeting cancer with small molecule kinase inhibitors. Nat. Rev. Cancer 9, 28–39.

52. Zhang, J., Cunningham, J.J., Brown, J.S., and Gatenby, R.A. (2017). Integrating evolutionary dynamics into treatment of metastatic castrate-resistant prostate cancer. Nat Comms 8, 1816.

53. Zhao, B., Hemann, M.T., and Lauffenburger, D.A. (2016a). Modeling Tumor Clonal Evolution for Drug Combinations Design. Trends in Cancer 2, 144–158.

54. Zhao, B., Sedlak, J.C., Srinivas, R., Creixell, P., Pritchard, J.R., Tidor, B., Lauffenburger, D.A., and Hemann, M.T. (2016b). Exploiting Temporal Collateral Sensitivity in Tumor Clonal Evolution. Cell 0.

