## Supplementary Figures for "Exploiting evolutionary herding to control drug resistance in cancer"

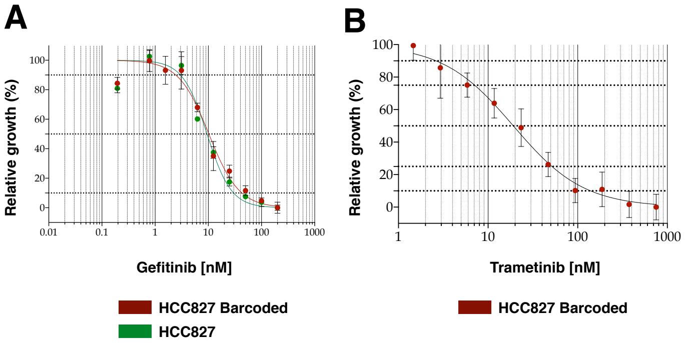


**Figure S1. Dose response curves of gefitinib and trametinib at baseline. (A)** Gefitinib and **(B)** trametinib dose response curves for baseline HCC827 used to estimate GI99 values for each drug.

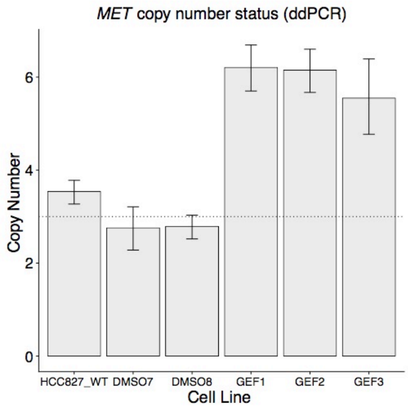


**Figure S2. MET amplification in gefitinib resistant lines is confirmed by ddPCR.** Number of copies of MET go from 3 in the baseline and DMSO to 6+ in the gefitinib evolved lines.

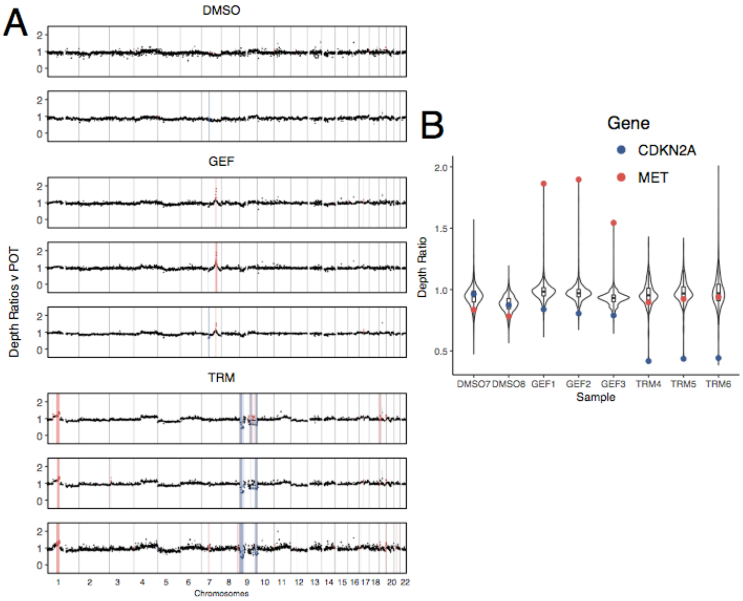


**Figure S3. Differential copy number profiles vs baseline POT. (A)** Depth ratio values of samples versus POT indicate that MET amplification in GEF lines and +1p and -9p in TRM lines. **(B)** Depth ratio values of MET and CDKN2A compare with whole-genome depth ratios highlight the putative driver changes in each line.

**Figure S4. Complete copy number calling per sample.** Each sample absolute copy number estimates are represented. HCC827 is a largely triploid cell line with large proportion of the genome being in LOH. This analysis confirmed the gain of 1p and 9q in TRM lines and the amplification of MET in the GEF lines.

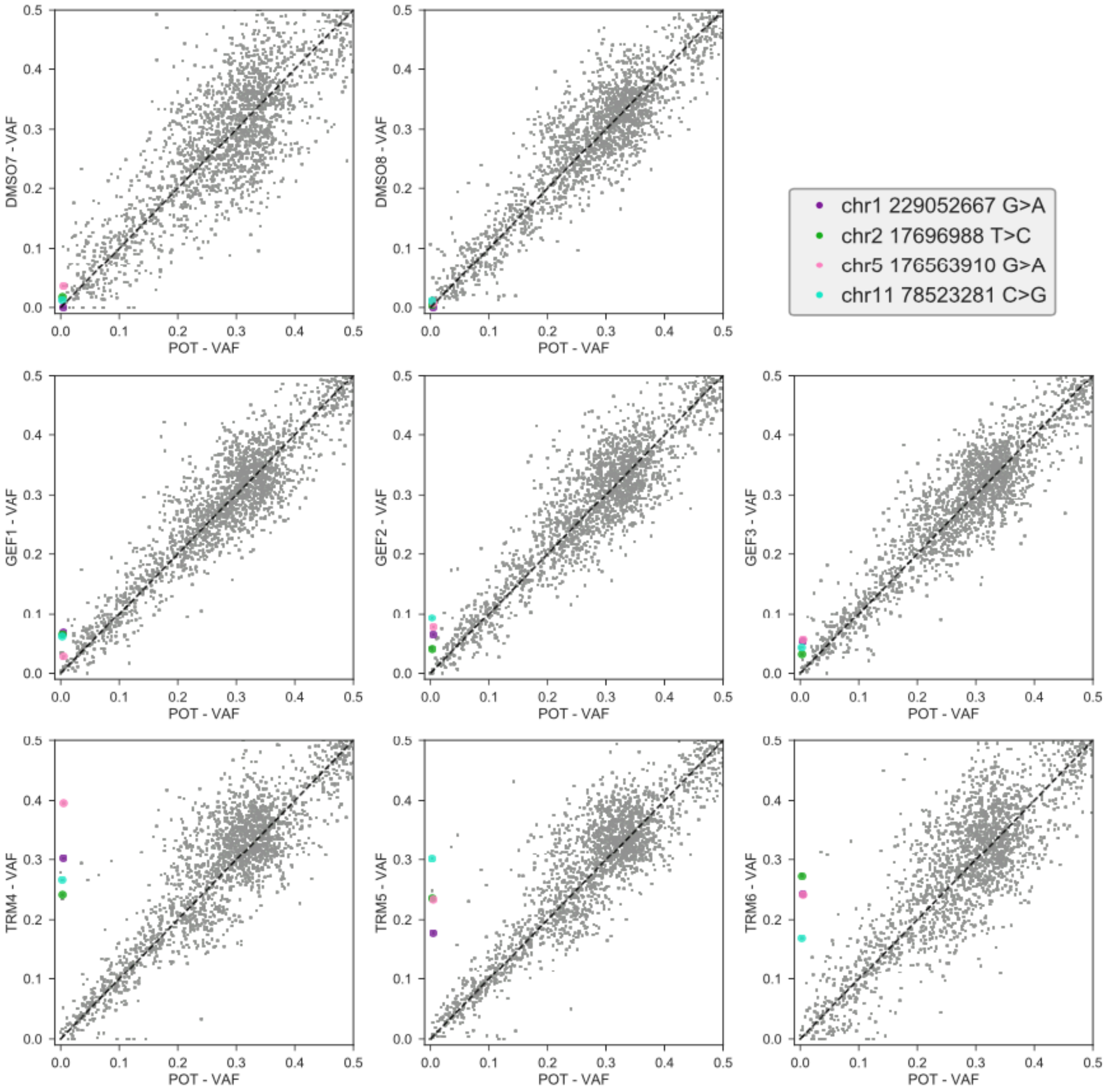


**Figure S5. SNV enrichment analysis.** HCC827 is a triploid cell line, hence we expect cluster of variants at VAF=0.33 and 0.66 (not shown). Subclonal mutations in a single allele that become clonal will reach clonality at VAF~0.33. A small set of mutations were enriched in all replicas of GEF and TRM, and reached almost clonality in TRM.

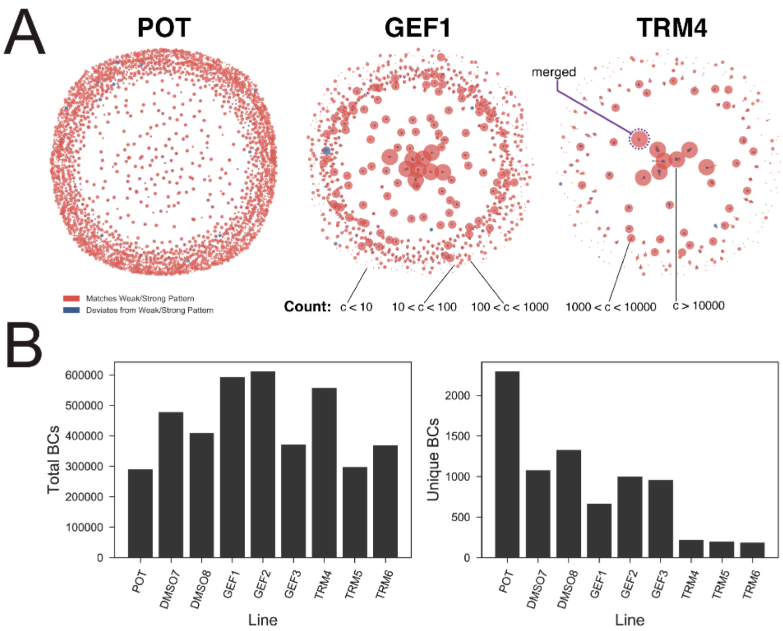


**Figure S6. Merging barcode sequences to correct for errors. (A)** Example barcode sequence distance networks for POT, GEF1 and TRM4. Points represent unique barcode sequences and are connected by edges where the sequences differ by Hamming distance 2 or less. Point size is determined by the number of the unique barcode detected in the replicate. Colour indicates whether the barcode matches the weak/strong basepair pattern. Our correction algorithm merges connected components of the network (example highlighted in purple). **(B)** Total barcodes (left) and unique barcodes (right) identified in each replicate following filtering and merging.

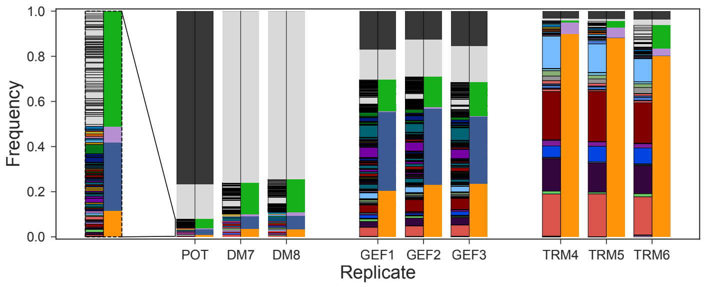


**Figure S7. Barcodes with undetermined phenotype.** Barcode frequency distributions in each sample as in Figure 4B. Left hand bars show the frequency of each unique barcode. Barcode colours are ordering are identical between replicates. Right hand bars indicate the phenotypes assigned to each barcode. Here, barcodes with undetermined phenotype (those not found in DM7 or DM8) are coloured dark grey.

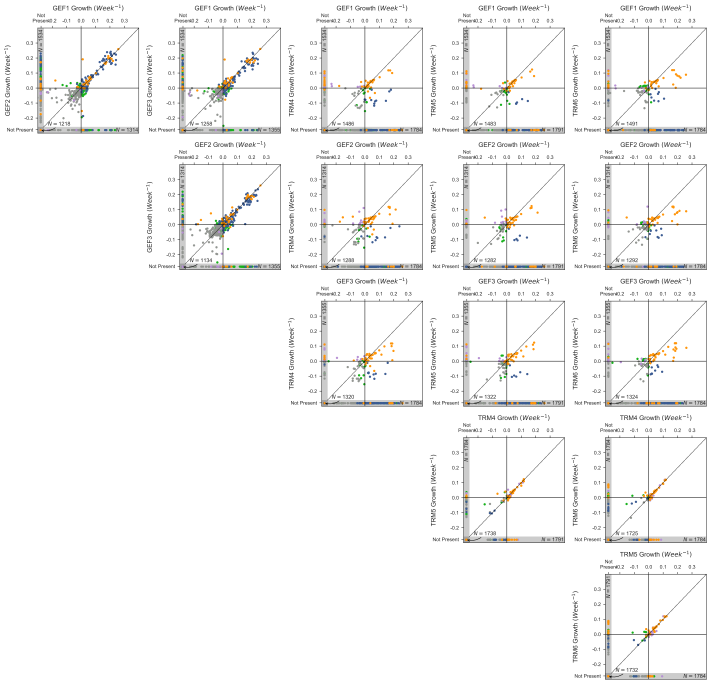

**Figure S8. Concordance in barcode growth rates between replicates.** Scatter plots show the concordance in barcode growth rates between pairs of evolutionary replicates. Points are coloured according to barcode phenotype, as in Figure 4B.

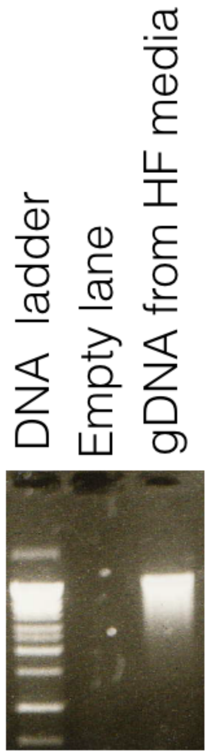


**Figure S9. DNA degradation demonstrate media floating cells are dead.** DNA from floating cells in the supernatant media was degraded, consistently with DNA coming from dead cells.

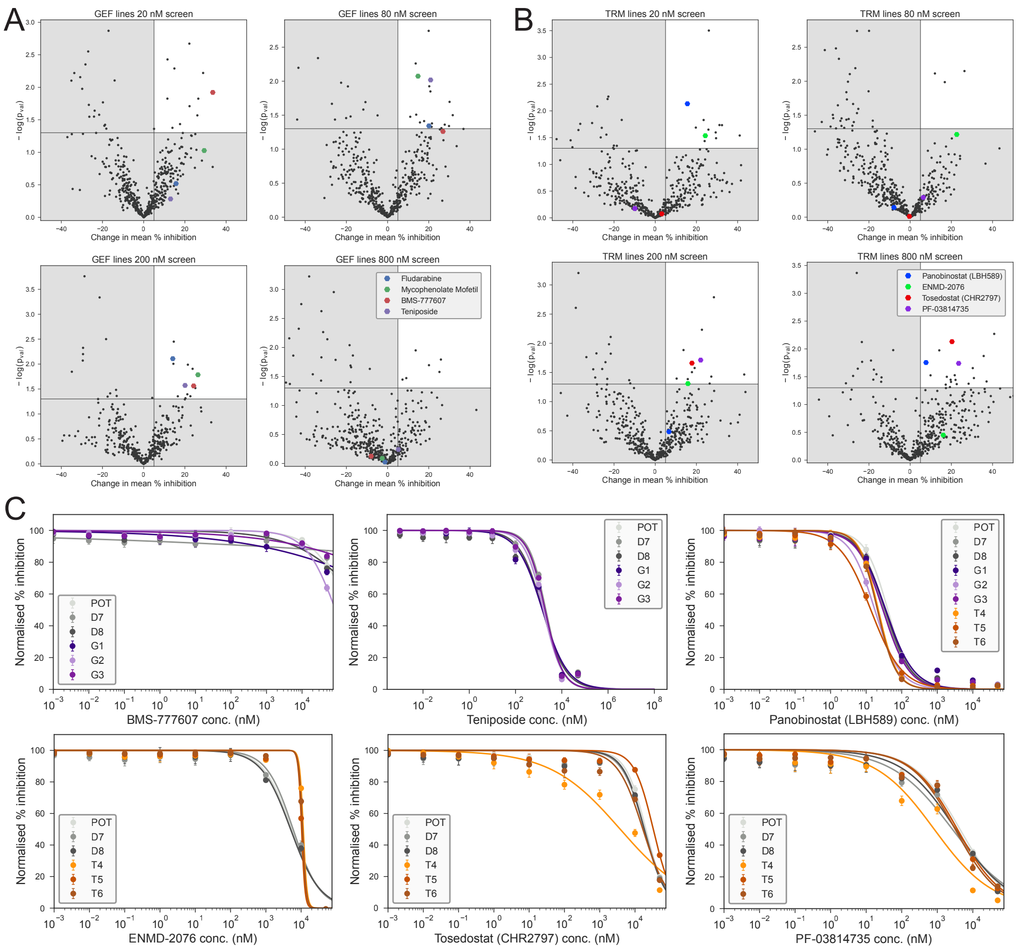


**Figure S10. High throughput screening for collaterally sensitive drugs. (A, B)** Volcano plots show the change in PCI and associated p-value for each compound at each concentration for the GEF, TRM lines. The upper quadrant delineates hits for collateral sensitivity. The 8 compounds that are hits are two or more concentrations are marked. **(C)** Dose response curves from the validation experiment for 6 of these compounds indicate no collateral sensitivity.

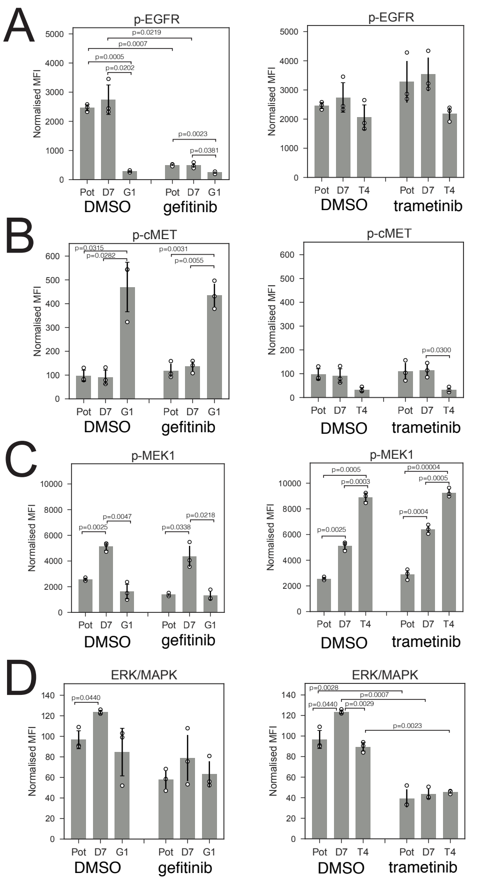


**Figure S11. Phosphoproteomics confirms drug action on signalling pathways. (A)** EGFR phosphorilation was highly downregulated under gefitinib and even in the absence of drug in GEF evolved lines, suggesting a stable phenotype where EGFR signalling has been lost due to clonal evolution. **(B)** MET phisphorilation was upregulated only in MET amplified GEF lines, as expected. **(C)** MEK phorphorilation was variable, however we confirmed ERK/MAPK downregulation under trametinib **(D)**, a strong indicator that the drug is inhibiting the MEK pathway. Error bars are determined by standard deviation, p-values are determined via a two-sided Welch's t-test.

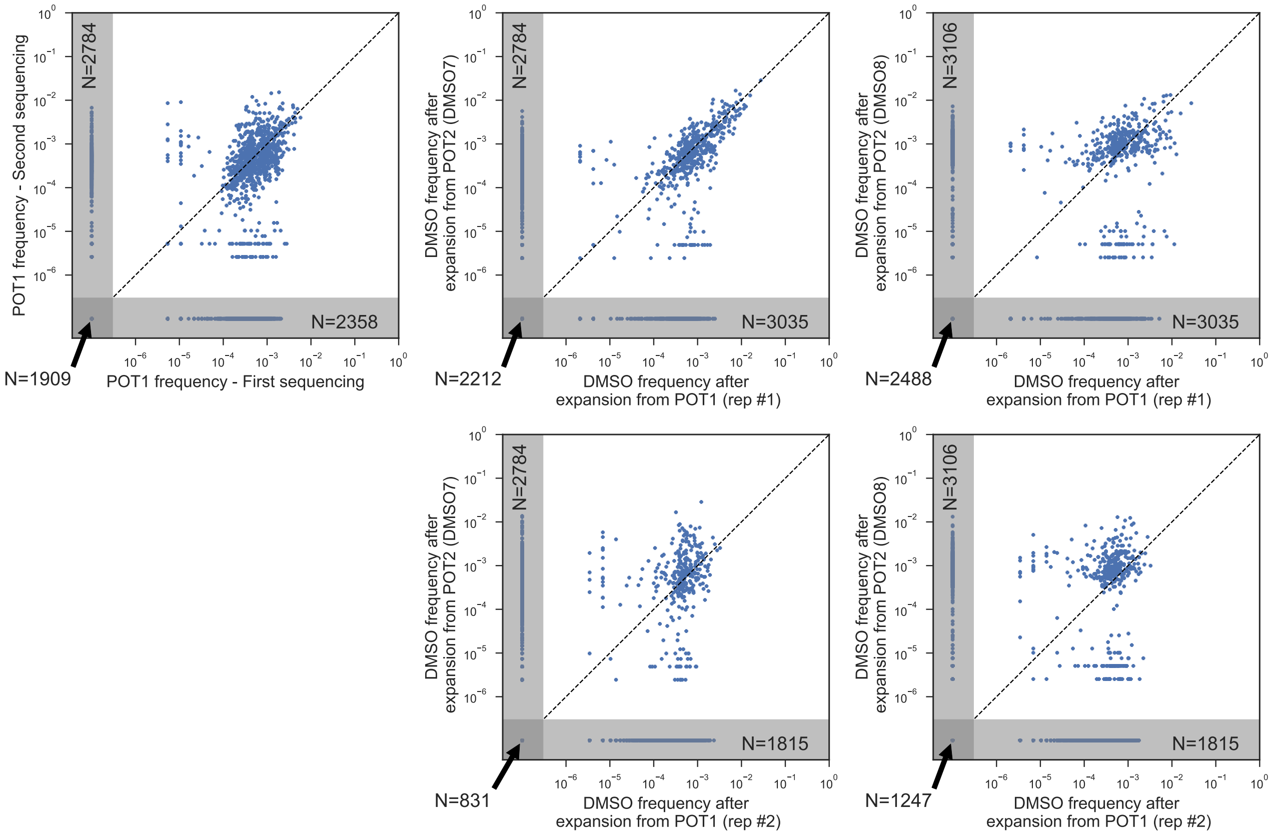


**Figure S12. Effects of freezing and thawing on barcode frequencies.** The frequencies of all barcodes identified in DMSO7 and DMSO8 are consistent to the frequencies in replicates expanded from the POT population before it was frozen. Some barcodes are always missed due to sequencing (binomial sampling of alleles).
