## Supplementary Notes for "Exploiting evolutionary herding to control drug resistance in cancer"

### Supplementary Methods

#### Statistical Analysis of Lentiviral Barcoding

##### Barcoding in HYPERFlasks

During the barcoding protocol, cells are randomly infected with barcodes such that one cell may receive multiple barcodes and that one barcode may appear in multiple cells. Following the statistical approach outlined by Lan et al. [2], we estimated the expected proportion of doubly barcoded cells and the expected proportion of barcodes that appear in multiple cells.

A brief overview of the approach of Lan et al. [2] proceeds as follows. Denote by $N_b$ the total number of barcodes, $p_b$ the probability of a barcoding event, and $n_b$ the number of barcodes received by a single cell. Then $n_b \sim \text{Poisson}(\nu)$ where $\nu = N_b p_b$. Thus,

$$\begin{aligned}
P_0 &= \mathbb{P}(n_b = 0) = e^{-\nu} \\
P_1 &= \mathbb{P}(n_b = 1) = \nu e^{-\nu} \\
P_{>1} &= \mathbb{P}(n_b > 1) = 1 - (1+\nu)e^{-\nu}.
\end{aligned}$$

$\nu$ can be estimated from the barcoding efficiency, $\eta$, defined as a the proportion of cells that are successfully labelled with at least one barcode. Specifically $\nu = -\log(1-\eta)$.

We estimate $\eta = 0.1$ as the proportion of cells which survive following selection with puromycin. Thus,

$$\begin{aligned}
P_0 &= 0.9 \\
P_1 &= 0.095 \\
P_{>1} &= 0.005.
\end{aligned}$$

Now, denote by $N_c$ the total number of cells prepared for lentiviral barcoding, and let $n_c$ be the number of cells receiving a specific barcode. Then $n_c \sim \text{Poisson}(\kappa)$ where $\kappa = p_b N_c = \frac{\nu N_c}{N_b}$. Thus,

$$\begin{aligned}
R_0 &= \mathbb{P}(n_c = 0) = e^{-\kappa} \\
R_1 &= \mathbb{P}(n_c = 1) = \nu e^{-\kappa} \\
R_{>1} &= \mathbb{P}(n_c > 1) = 1 - (1+\kappa)e^{-\kappa}.
\end{aligned}$$

Hyo-eun et al. [1] report a total barcode library complexity $N_b = 7.2 \times 10^7$ by fitting a polynomial equation. The total number of cells prepared for barcoding was $N_c = 10^7$. Thus,

$$R_0 = 0.986$$
$$R_1 = 0.014 \qquad\qquad R_{>1} < 0.001.$$

Finally, the proportion of uniquely labelled cells (those that receive a unique combination of one or more barcodes) is given by

$$Q \approx \frac{1 - P_0^{R_0}}{1 - P_0^{-1}} = 0.985.$$

##### Barcoding in Organoids

For organoid barcoding we estimate $N_c = 9 \times 10^6$, $\eta = 0.1$ and the barcode library complexity is as above. We estimate

$$P_0 = 0.9$$
$$P_1 = 0.095$$
$$P_{>1} = 0.005$$
$$R_0 = 0.987$$
$$R_1 = 0.013$$
$$R_{>1} < 0.001$$

and $Q \approx 0.986$.

#### Barcode Sequence Merging

##### Barcodes from Harvested Cells

Errors introduced during PCR or sequencing of the molecular barcodes can result in spurious barcodes being identified, or in the underestimation of the prevalence of a specific barcode. We implemented a novel, bias free error correction protocol as follows.

All reads matching the 12bp of the forward primer, followed by a 30bp sequence, followed by 12bp of the reverse primers were considered. This permits us to identify barcodes that deviate from the weak/strong base pair pattern as a result of errors. Reads were filtered for base quality score >20 in all positions. All detected barcodes were merged into a single file to ensure that the same corrections were applied between different samples.

We next constructed a graph in which the vertices are unique barcodes and the edges connect barcodes which differ by Hamming distance at most 2. We propose that the connected components of this graph represent groups of barcode sequences derived from the same true barcode, and thus that connected components (CCs) should be merged into single representative barcodes. Figure S6 shows the Hamming distance networks for POT, GEF1 and TRM4. A total of 4777 connected components were extracted. Connected components not containing a representative barcode matching the weak/strong barcode pattern were discarded (60 CCs containing 103 unique barcodes and 12914 total post-filtering reads, 0.0032% of all of the post–filtering reads).

A representative barcode was selected from each CC as the most abundant barcode matching the the weak/strong pattern. Where this representative was >10× more abundant than every other barcode in the CC, all others were corrected to the representative. Where there existed additional barcodes matching the weak/strong pattern and >0.1× the representative count, we designated these barcodes as alternative representatives. 46 CCs were identified as containing multiple representatives (ranging from 2-5 representatives). For these CCs the barcodes were split amongst the representatives by correcting each barcode to the representative with the smallest Hamming distance. Where multiple representatives had the same Hamming distance, the count for the barcode was evenly split between the representatives to avoid bias.

**Barcodes from Supernatent Cells**

For the barcodes extracted weekly from the supernatent cells, the extraction and filtering were performed as above. The correction mapping derived from harvested cells was used to correct the barcodes. There were no barcodes identified in the supernatent cells that was not detected in at least one of the harvested samples.

### References


1. C Bhang Hyo-eun, David A Ruddy, Viveksagar Krishnamurthy Radhakrishna, Justina X Caushi, Rui Zhao, Matthew M Hims, Angad P Singh, Iris Kao, Daniel Rakiec, Pamela Shaw, et al. Studying clonal dynamics in response to cancer therapy using high-complexity barcoding. *Nature medicine*, 21(5):440, 2015.

2. Xiaoyang Lan, David J Jörg, Florence MG Cavalli, Laura M Richards, Long V Nguyen, Robert J Vanner, Paul Guilhamon, Lilian Lee, Michelle M Kushida, Davide Pellacani, et al. Fate mapping of human glioblastoma reveals an invariant stem cell hierarchy. *Nature*, 549 (7671):227, 2017.

3. Shannon M Mumenthaler, Jasmine Foo, Nathan C Choi, Nicholas Heise, Kevin Leder, David B Agus, William Pao, Franziska Michor, and Parag Mallick. The impact of mi-

croenvironmental heterogeneity on the evolution of drug resistance in cancer cells. *Cancer informatics*, 14:CIN–S19338, 2015.
